# A Potential Role for Aminoacylation in Primordial RNA Copying Chemistry

**DOI:** 10.1101/2020.12.06.413815

**Authors:** Aleksandar Radakovic, Tom H. Wright, Victor S. Lelyveld, Jack W. Szostak

## Abstract

Aminoacylated tRNAs are the substrates for ribosomal protein synthesis in all branches of life, implying an ancient origin for aminoacylation chemistry. In the 1970s, Orgel and colleagues reported potentially prebiotic routes to aminoacylated nucleotides and their RNA templated condensation to form amino acid bridged dinucleotides. However, it is unclear whether such reactions would have aided or impeded nonenzymatic RNA replication. Determining whether aminoacylated RNAs could have been advantageous in evolution prior to the emergence of protein synthesis remains a key challenge. We therefore tested the ability of aminoacylated RNA to participate in both templated primer extension and ligation reactions. We find that at low magnesium concentrations that favor fatty acid-based protocells, these reactions proceed orders of magnitude more rapidly than when initiated from the cis-diol of unmodified RNA. We further demonstrate that amino acid bridged RNAs can act as templates in a subsequent round of copying. Our results suggest that aminoacylation facilitated nonenzymatic RNA replication, thus outlining a potentially primordial functional link between aminoacylation chemistry and RNA replication.

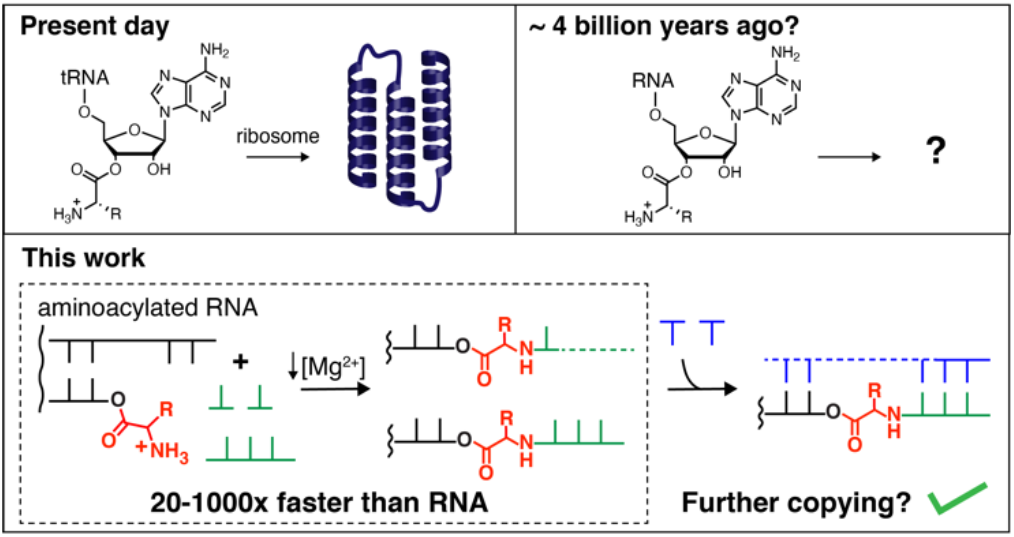

## INTRODUCTION

In extant biochemistry, the aminoacylation of RNA generates activated tRNA substrates for protein biosynthesis. Explaining how RNA could be aminoacylated without enzymes, and how such aminoacylation might have benefitted early protocells, may help to explain how the RNA World transitioned to protein-centric biology. The covalent linkage of amino acids to RNA would have unavoidably affected RNA replication. If these effects were beneficial, then efficient ribozyme catalyzed aminoacylation could have evolved. Once in place, ribozyme catalysis of aminoacylation could in turn have led to other uses for covalently attached amino acids, such as peptide formation.

Nucleotides and RNA strands can be aminoacylated at the 2’(3’) hydroxyl groups by reaction with amino acid imidazolides^1^ (Figure 1A), which in turn can be formed from the imidazole-catalyzed reaction of a free amino acid with a nucleotide activated as a 5’-phosphorimidazolide^2^. However, the latter reaction competes with the rapid conversion of the aminoacyl adenylate intermediate into an N-carboxyanhydride (NCA) in the presence of CO_2_^3, 4^. NCAs are inefficient reagents for the direct aminoacylation of ribonucleotides^5^. High yielding aminoacylation pathways employing NCAs^6^ or in situ activation chemistry^7^ are known, but they require a 3’-phosphate moiety, and are generally limited to N-blocked amino acids. Therefore, we still do not have a high yielding and prebiotically plausible means to chemically aminoacylate RNA strands terminating in a vicinal diol. However, even inefficient chemistry could have had significant effects on RNA replication and assembly processes.

**Figure 1.**
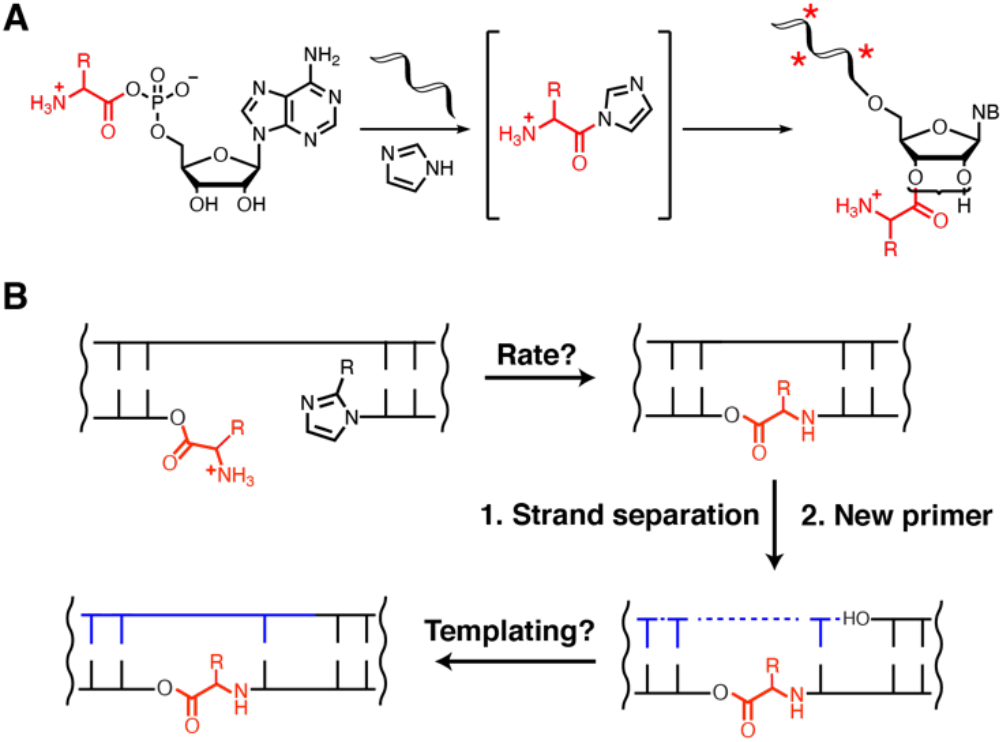
Aminoacylation of RNA and integration of covalently attached amino acids with RNA copying. **(A)** Aminoacylation chemistry. The reaction of imidazole with aminoacyl adenylate anhydrides generates amino acid imidazolides, which in turn serve as amino acid donors for covalent modification of RNA strands with amino acids (asterisks are internal 2’ acylated OHs). **(B)** Nonenzymatic primer extension or ligation initiated from a 2’(3’) aminoacyl terminated RNA strand generates amino acid bridged RNA, which can act as a template for future rounds of copying.

We were curious as to whether the additional chemical functionality imparted by amino acids could assist the non-enzymatic, chemical copying of RNA strands. In 1974, Shim and Orgel reported that 2’(3’)-aminoacylated nucleotides can react with nucleotide 5’-phosphorimidazolides in a template-directed process to form phosphoramidate-linked products via transamidation^8^. On a poly(U) template, 2’(3’)-glycyl adenosine reacted with 5’-imidazole activated AMP to afford a dinucleotide species that was bridged by the glycine residue^8^. Formation of such phosphoramidate linked amino-acid RNA ‘co-polymers’ dramatically increases the stability of both the aminoacyl ester and phosphoramidate bonds, suggesting a mechanism for enhanced covalent capture of amino acids by early RNA^9^. In parallel, the Orgel laboratory^10,11^ and later our own group^12–14^ demonstrated that nucleotides with either 2’- or 3’-amino groups exhibit a large increase in the rate and yield of non-enzymatic polymerization due to the greater nucleophilicity of the amine substituent relative to a hydroxyl group. These results have been extended to ligation reactions by the Krishnamurthy group^15^ and our own recent assembly of an active ribozyme via a series of 3’N-5’P ligation reactions^16,17^. Together, these results inspired us to test whether the reaction of the free amino group of 2’(3’) aminoacylated RNA with phosphorimidazolide activated nucleotides would also result in enhanced rates of RNA copying, and whether such amino acid-bridged oligonucleotides could act as templates for cycles of replication (Figure 1B).

Here, we report that terminal 2’(3’) aminoacylation of RNA can indeed enhance both template-directed primer extension and ligation reactions with imidazole activated downstream species by over two orders of magnitude. We also show that RNA oligomers containing a single amino acid ‘bridge’ can act as templates for subsequent rounds of RNA template copying. Taken together, our findings reveal a potential pathway by which amino acids may have initially become integrated into the RNA world, thus setting the stage for the subsequent emergence of RNA-encoded protein synthesis.

## MATERIALS AND METHODS

### General information

All reagents were purchased from Sigma-Aldrich (St. Louis, MO) unless specified otherwise. TurboDNase was purchased from Thermo Scientific (Waltham, MA). Flexizyme “dFx” and the corresponding mutant M2 were prepared as described elsewhere^18^. PCR was performed with Hot Start Taq 2X Master Mix and in-vitro transcription with HiScribe™ T7 Quick High Yield RNA Synthesis Kit from New England Biolabs (Ipswich, MA). EDTA is used as an abbreviation for Na_2_EDTA pH 8.0.

### Oligonucleotide synthesis

Oligonucleotides were either purchased from Integrated DNA Technologies (Coralville, IA) or synthesized in house on an Expedite 8909 solid-phase oligo synthesizer. Phosphoramidites and reagents for the Expedite synthesizer were purchased from either Glen Research (Sterling, VA) or Chemgenes (Wilmington, MA). Cleavage of synthesized oligonucleotides from the solid support was performed using 1 mL of AMA (1:1 mixture of 28 % aqueous ammonium hydroxide and 40 % aqueous methylamine) for 30 minutes at room temperature, while deprotection was done in the same solution for 20 minutes at 65 °C. Deprotected oligos were lyophilized, resuspended in 100 μL DMSO and 125 μL TEA.3HF, and heated at 65 °C for 2.5 h to remove TBDMS from 2’ hydroxyls. Following this deprotection, oligos were purified by preparative 20 % polyacrylamide gel electrophoresis (19:1 with 7 M Urea), desalted using Waters Sep-Pak C18 cartridges (Milford, MA), and characterized by high-resolution mass spectrometry on an Agilent 6230 TOF mass spectrometer.

### Oligonucleotide and nucleotide activation

Oligonucleotides phosphorylated on the 5’ OH were activated with 2-methylimidazole or 2-aminoimidazole as previously reported^19^ with the following modifications: gel-purified products of 1 μmol solid-phase synthesis were dissolved in 100 μL DMSO. 0.05 mmol of triethylamine (TEA), 0.02 mmol of triphenylphosphine (TPP), 0.04 mmol of 2-methylimidazole (or 2-aminoimidazole), and 0.02 mmol of 2,2’-dipyridyldisulfide (DPDS) were added to the reaction, and the reaction was incubated on a rotator for 5 h at room temperature. After 5 h, all of the reagents above were added in listed quantities again and the reaction was allowed to rotate for an additional 12 h at room temperature. The reaction was precipitated with 100 μL saturated NaClO_4_ in acetone and 1 mL acetone for 30 minutes on dry ice. The pellet was washed with 1 mL 1:1 acetone:diethylether twice. The products were resolved and purified by HPLC on an Agilent ZORBAX analytical column (Eclipse Plus C18, 250 × 4.6mm, 5 μm particle size, P.N. 959990-902), at a flow rate of 1 ml/min. Gradient: (A) aqueous 20 mM triethylammonium bicarbonate pH 8.0, and (B) acetonitrile, from 7 % to 12 % B over 12 minutes.

2-aminoimidazolium cytidine dinucleotide (C*C): 0.46 mmol of CMP (free acid) was dissolved in 4 mL DMSO. 2.9 mmol of TEA, 3.8 mmol of TPP, and 0.22 mmol of 2-aminoimidazole (HCl salt) were added to the CMP solution. The resulting suspension was sonicated and heated briefly until all reagents completely dissolved. 4 mmol of DPDS was added to the solution to start the reaction and the reaction was stirred for 15 minutes at room temperature. The reaction was then precipitated by adding 0.5 mL of saturated NaClO_4_ in acetone and 60 mL 1:1 acetone:diethyl ether, washed with 1:1 acetone:diethyl ether twice, and purified by C_18_ reverse-phase chromatography at a 40 mL/min flow rate. Gradient: (A) aqueous 2 mM triethylammonium bicarbonate pH 8.0, and (B) acetonitrile, from 0 % to 10 % B over 10 minutes.

2-aminoimidazolium guanosine-uridine dinucleotide (G*U): OAt-GMP synthesis: 0.275 mmol of GMP (free acid) was dissolved in 18 mL of water, followed by 1.47 mmol of HOAt and 1.5 mmol of TEA. The solution was then lyophilized. The resulting powder was dissolved in 10 mL DMSO, followed by the addition of 3.6 mmol of TEA, 2.75 mmol of TPP, and 2.75 mmol of DPDS. The reaction was stirred at room temperature for 30 mins, precipitated by adding 0.5 mL of saturated NaClO_4_ in acetone and 60 mL 1:1 acetone:diethyl ether, washed with 1:1 acetone:diethyl ether twice, and purified by C_18_ reverse-phase chromatography at a 40 mL/min flow rate. Gradient: (A) aqueous 2 mM triethylammonium bicarbonate pH 8.0, and (B) acetonitrile, from 0 % to 15 % B over 10 minutes. 2-AI-UMP synthesis: UMP was activated as C*C, except 2-aminoimidazole was used at 3 eq. The purified OAt-GMP and 2-AI-UMP were then mixed in 4 mL water for 1 hour. Purification of G*U was performed by preparative HPLC on an Agilent preparative column (Eclipse XDB C_18_, 250 × 21.2 mm, 7 μm particle size, P.N. 977250-402), at a flow rate of 15 ml/min. Gradient: (A) aqueous 2 mM triethylammonium bicarbonate pH 8.0, and (B) acetonitrile, from 2 % to 8 % B over 20 column volumes.

### Amino acid substrate synthesis

Amino acid-DBE substrates (3,5-dinitrobenzyl esters of amino acids) were synthesized as reported previously^18^ with the following modification: *N*-Boc protected amino acid DBE-esters were deprotected in 2 mL neat TFA for 10 minutes, followed by washing with 3x 10 mL diethyl ether. Products were obtained as TFA salts. TFA salts were dissolved in 100 % DMSO to 25 mM and used in reactions directly. ^1^H NMR spectra were obtained using a 400 MHz NMR spectrometer (Varian INOVA) operating at 400 MHz. Low resolution mass spectrometry was performed by directly injecting ~ 2 mg of material in 1 mL 1:1 acetonitrile:water on an Esquire 6000 mass spectrometer (Bruker Daltonics). High resolution mass spectrometry was performed by injecting 500 pmol of material dissolved in water on an Agilent 1200 HPLC coupled to an Agilent 6230 TOF mass spectrometer.

### Flexizyme catalyzed aminoacylation of oligonucleotides (Figure S1)

Aminoacylation reactions were performed as reported^18^ with the following modifications: a typical 10 μL reaction contained 50 mM Na-HEPES pH 8.0, 10 mM MgCl_2_. 10 μM fluorescein-labeled primer, 5 mM aa-DBE (20 % DMSO final), 10 μM dFx Flexizyme. The reaction was incubated on ice for 12-16 h. 1 μL of the reaction was quenched with 9 μL quench buffer (10 mM EDTA, 100 mM NaOAc pH 5.0, 150 mM HCl, 70 % v/v formamide) and loaded into 20 % polyacrylamide gel (19:1 with 7 M Urea, 0.1 M NaOAc pH 5.0) in the cold room (4 °C). The gel was run for 2 h at 300 V and visualized on a Typhoon 9410 imager. A typical aminoacylation reaction yielded 30-60 % product, measured by band densities in ImageQuant TL software.

### Amino acid-bridged oligonucleotide synthesis (Figure S2)

Primer with the amino acid bridge before the 3’ terminal nucleotide (**1**) was synthesized as described previously^20^ with the following modifications: oligonucleotide aminoacylation was performed at a 1 mL scale, then split into two followed by the addition of the FX_T2 hybrid template and FX_S2 “sandwich” (Table S1) to the final concentration of 2.5 μM each. Na-HEPES pH 8.0 and EDTA were added to final concentrations of 200 mM and 50 mM, respectively. The solution was allowed to warm to room temperature for 2 minutes, after which the reaction was started by the addition of the C*C dinucleotide to the final concentration of 13.5 mM. The reaction was allowed to proceed for 10 minutes while rotating at room temperature. The reaction was then concentrated using Amicon Ultra-4 mL 3K centrifugal filters and the buffer was exchanged twice with nuclease-free water. The reactions were combined and further concentrated down to 50 μL with Amicon Ultra-0.5 mL 3K centrifugal filters. 375 μL of nuclease-free water, 50 μL of 10x TurboDNase buffer, and 50 U of 2 U/μL TurboDNase were added. TurboDNase digestion was allowed to proceed for 15 minutes at 37 °C. The digested reaction was then concentrated using Amicon Ultra-0.5 mL 3K centrifugal filters, diluted with 5 mM EDTA in 95 % v/v formamide, and purified by preparative 20 % polyacrylamide gel electrophoresis (19:1 with 7 M Urea) at 4 °C. The desired gel band was cut out, crushed, and extracted with 1 mL of 50 mM NaOAc pH 5.5 50 mM EDTA buffer on a rotator at 4 °C for 16 h. The extracted product was concentrated using Amicon Ultra-0.5 mL 3K centrifugal filters and the buffer was exchanged with nuclease-free water three times to yield 80-90 % pure **1**.

Template with an internal amino acid bridge (**2**) was synthesized by performing a ligation reaction with the following modifications: Oligonucleotide aminoacylation was performed at a 1 mL scale, then split into two followed by the addition of the NP DNA T template (Table S1) to the final concentration of 2.5 μM. Na-HEPES pH 8.0 and EDTA were added to final concentrations of 200 mM and 50 mM, respectively. The solution was allowed to warm to room temperature for 2 minutes, after which the reaction was started by the addition of 2-methylimidazole activated Ligator1 (Table S1) to the final concentration of 10 μM. The reaction was allowed to proceed for 1 h while rotating at room temperature. The reaction was concentrated with Amicon Ultra-4 mL 3K centrifugal filters and the buffer was exchanged with nuclease-free water twice. The reaction was further concentrated to 50 μL using Amicon Ultra-0.5 mL 3K centrifugal filters. DNA digestion and purification were performed exactly as for **1**. After gel extraction and buffer exchange, the 90 % pure **2** was precipitated with 0.1 volume of 5 M NH_4_OAc and 3 volumes of isopropanol.

### Primer extension reactions

With C*C dinucleotide (Figures 2, 3, 4, S5, and S6):

RNA template and the downstream RNA oligonucleotide (“sandwich”) were added to a typical 10 μL aminoacylation reaction to the final concentrations of 3.75 and 2.5 μM, respectively, followed by Na-HEPES pH 8.0 to a final concentration of 200 mM. MgCl_2_ was added to a final concentration of 50 mM for reactions that were performed at 50 mM MgCl_2_; water was added to reactions performed at 2.5 mM MgCl_2_; EDTA was added to reactions performed at 0 mM MgCl_2_ to a final concentration of 25 mM. Note: because the aminoacylation reactions were performed in the presence of 10 mM MgCl_2_, they contributed 2.5 mM of MgCl_2_ to the final reaction. The reactions were allowed to warm up to room temperature for 2 minutes before being initiated by the addition of the C*C dimer to a final concentration of 20 mM. Final reaction concentrations: 2.5 μM mixture of primers, 0-50 mM MgCl_2_, 200 mM HEPES pH 8.0, and 20 mM C*C. Reactions were performed in technical triplicates. At indicated time points, 1 μL of each reaction was quenched with 29 μL quench buffer (final quench buffer concentrations: 50 mM EDTA, 2 μM reverse complement of the template, 90 % v/v formamide). Prior to loading on 20 % polyacrylamide gels (19:1 with 7 M Urea), the quenched reactions were heated at 92 °C for 2 minutes to denature the duplex. 3 μL aliquots were loaded into gels and run at 20 W for 1 h 20 minutes. The gels were imaged on Typhoon 9410 imager and band densities quantified in ImageQuant TL software.

**Figure 2.**
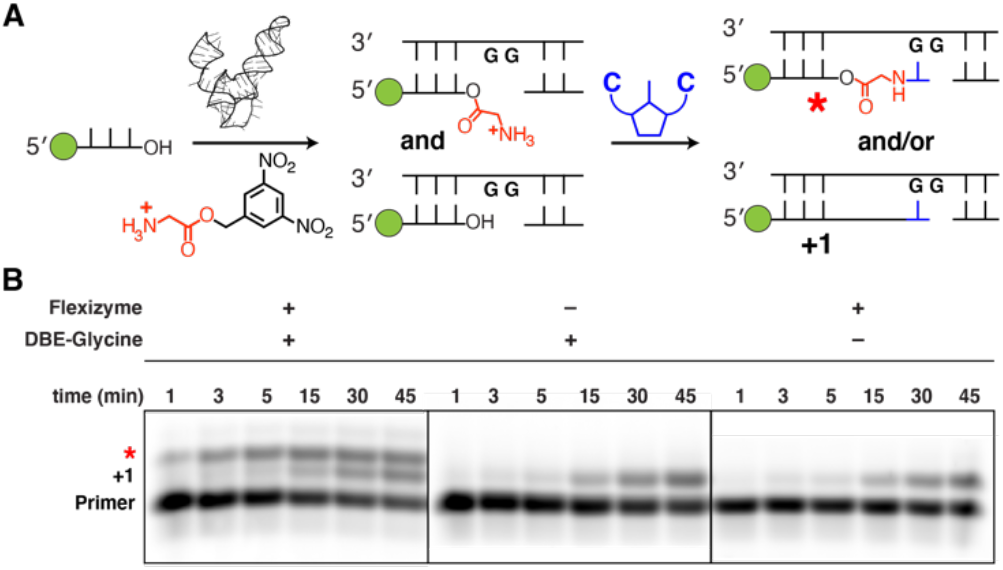
Nonenzymatic copying of RNA initiated from 2’(3’) aminoacyl terminated RNA. **(A)** Schematic of the assay used. Flexizyme acylation of a fluorescently-labeled (green circle = fluorescein) RNA primer results in a mixture of acylated and nonacylated strands which are used in primer extension with the 5’-5’ aminoimidazolium-bridged cytidine dinucleotide, C*C. **(B)** The time course of primer extension was monitored using polyacrylamide gel electrophoresis. All reactions were performed at pH 8.0, 200 mM HEPES, 2.5 mM MgCl_2_, with 20 mM C*C.

**Figure 3.**
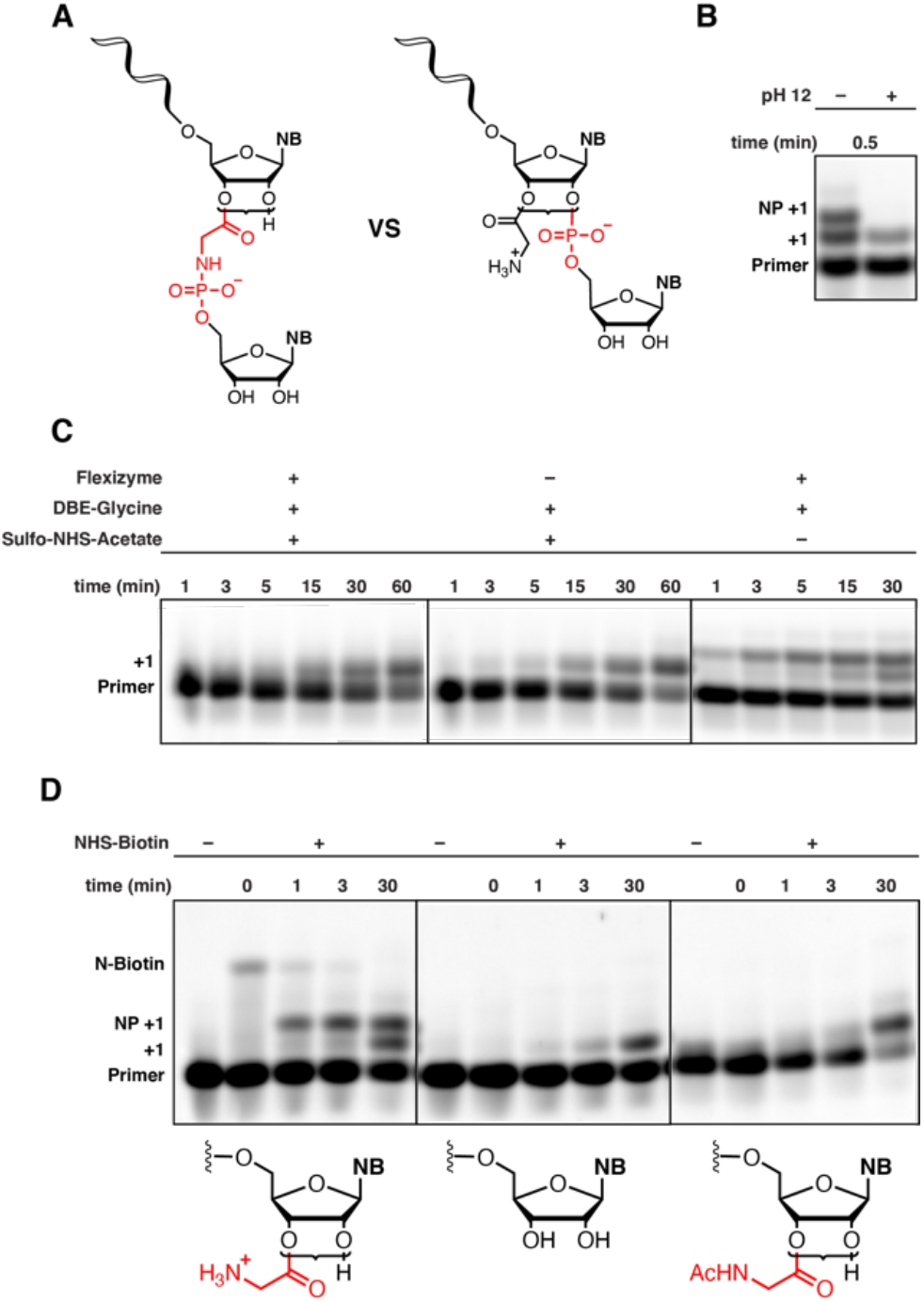
Confirmation of the presence of an amino acid ‘bridge’. **(A)** Possible reaction products of nonenzymatic copying initiated from 2’(3’) aminoacyl terminated RNA. Brackets indicate a dynamic mixture of 2’(3’) aminoacylated RNA. **(B)** Treatment with NaOH leads to disappearance of the novel ‘NP+1’ band, suggesting the presence of an aminoacyl ester. **(C)** Chemical N-acetylation prevents appearance of the novel ‘NP+1’ band, suggesting that the glycyl amino group is required for formation of the novel +1 product. **(D)** Formation of the phosphoramidate linked product inhibits reaction with NHS-biotin. As the primer is extended, the concentration of the glycyl amino group declines due to N-P bond formation, leading to reduced labelling with the biotinylation reagent. All reactions were performed at pH 8.0, 200 mM HEPES, 2.5 mM MgCl_2_, with 20 mM C*C dimer.

**Figure 4.**
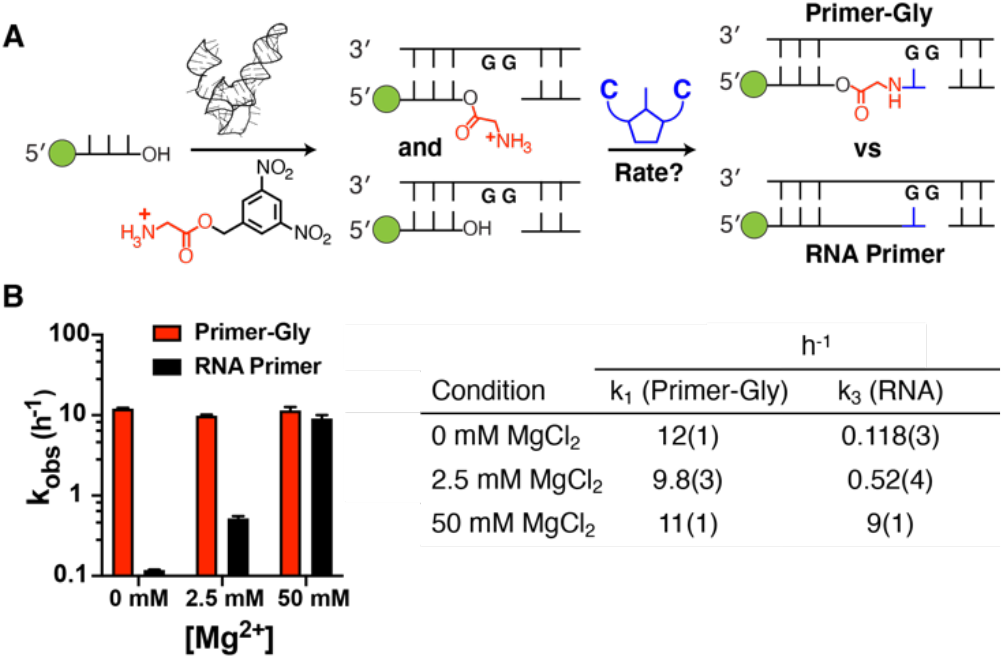
Kinetic analysis of nonenzymatic primer extension from 3’ hydroxyl and 2’(3’) glycyl terminated primers. **(A)** Schematic representation of the assay. **(B)** Kinetic parameters. Left: k_obs_ (h^-1^) vs. concentration of Mg^2+^. Right: pseudo first-order rate constant k_1_, for extension of the aminoacylated primer, and k_3_, the native RNA primer extension rate constant, at 0, 2.5 and 50 mM MgCl_2_. Rate constants were estimated by nonlinear regression using values for k_2_, the aminoacyl hydrolysis rate constant, and the initial fraction of aminoacylated primer Pgly0, that were estimated from independent measurements. Plots used to obtain k_1_, k_2_ and k_3_ can be found in Figure S5. Uncertainty in the estimate of k_1_ was analyzed by the Monte Carlo method to propagate error on the estimates of independent model parameters. All reactions were performed at pH 8.0, 200 mM HEPES, with 20 mM C*C dimer. Values are reported as the mean +/- standard deviation from triplicate experiments.

To independently determine the kinetics of pure RNA primer extension, primer extension was performed with the non-aminoacylated RNA primer. The non-aminoacylated primer was subjected to typical aminoacylation conditions, except dFx Flexizyme was replaced with water for those reactions. The primer extension reactions were then set up exactly as in the preceding paragraph.

The rate of hydrolysis of the aminoacylated primer was measured under primer extension conditions except that CMP was used instead of C*C. The hydrolysis reaction was quenched using the acidic quench buffer (10 mM EDTA, 100 mM NaOAc pH 5.0, 150 mM HCl, 2 μM reverse complement of the template, 70 % v/v formamide), heated at 92 °C for 2 minutes to denature the duplex, and run on acidic 20 % polyacrylamide gel (19:1 with 7 M Urea, 0.1 M NaOAc pH 5.0).

With C*C dinucleotide (Figure S7):

The purified product **1** and the downstream “sandwich” were annealed to the RNA template in a solution containing 3.6 μM **1**, 3.6 μM “sandwich”, 5.4 μM template, 50 mM Na-HEPES pH 7.5, 50 mM NaCl, and 1 mM EDTA by heating for 3 minutes at 70 °C and slowly cooling to 20 °C at 0.1 °C/s. The annealed solution was diluted with Na-HEPES pH 8.0 and MgCl_2_, before initiating the reaction by adding the C*C dinucleotide. The final reaction concentrations were 0.6 μM **1**, 200 mM Na-HEPES pH 8.0, 50 mM MgCl_2_, and 20 mM C*C. The reaction was quenched at indicated time points, subjected to gel electrophoresis, and quantified as described above.

With G*U dinucleotide (Figures 6 and S12):

The primers and the corresponding downstream “sandwich” oligonucleotides were annealed to the template (either the glycine-linked **2** or the all RNA template) in a solution containing 3.6 μM primer, 3.6 μM “sandwich”, 5 μM template, 50 mM Na-HEPES pH 7.5, 50 mM NaCl, and 1 mM EDTA by heating for 3 minutes at 70 °C and slowly cooling to 20 °C at 0.1 °C/s. The annealed solution was diluted with Na-HEPES pH 8.0 and MgCl_2_, before initiating the reaction by adding the G*U dinucleotide. The final reaction concentrations were 0.6 μM primer, 200 mM Na-HEPES pH 8.0, 100 mM MgCl_2_, and 20 mM G*U. The reaction was quenched at indicated time points, subjected to gel electrophoresis, and quantified as described above.

### Kinetic analysis of primer extension reactions

With C*C dinucleotide (Figures 2, 4, and S5):

RNA reaction: primer extension was quantified for each time point by integrating band intensity in each gel lane. Band intensity was normalized in each lane. Remaining primer (P) at each time point, starting from the initial fraction of primer (P_0_), was plotted as −ln(P/P_0_) vs. reaction time, and the observed rate constant, k_obs_, was estimated by the slope of a linear regression line. This k_obs_ corresponded to k_3_ in the kinetic model used to obtain k_obs_ (k_1_) of the aminoacylated primer.

Hydrolysis reaction: hydrolysis was quantified for each time point by integrating band intensity in each gel lane. Band intensity was normalized in each lane. Remaining primer-gly (P) at each time point, starting from the initial fraction of primer-gly (P_0_), was plotted as −ln(P/P_0_) vs. reaction time, and the observed rate constant, k_obs_, was estimated by the slope of a linear regression line. This k_obs_ corresponded to k_2_ in the kinetic model used to obtain k_obs_ (k_1_) of the aminoacylated primer.

Aminoacylated reaction: to obtain a rate constant for extension of the aminoacylated primer under conditions saturating for the 2-aminoimidazolium-bridged dinucleotide, we modeled the reaction with the following simplified kinetic scheme.

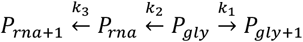

In this reaction network, *P_rna_* is the native RNA primer, which can be formed by hydrolysis from *P_gly_*, the glycyl-terminal primer. The respective +1 species are corresponding extended products of each form of the primer. The total observable primer concentration is *P* = *P_gly_* + *P_rna_*, and the normalized extent of initial aminoacylation of the primer is governed by the expression *P_rna_0__* = 1 – *P_gly_0__*. Under the assumption of pseudo-first-order kinetics, the reaction can be described by the following system of differential equations.

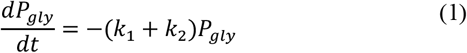

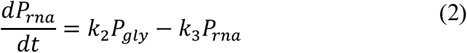

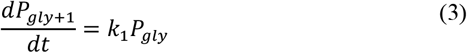

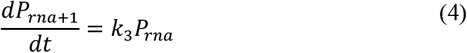

Integrating (1) yields an expression for the consumption of glycyl-terminal primer.

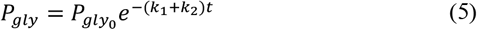

The differential equation (3) can be integrated with (5), yielding an expression for the extension of glycyl-terminal primer to form the +1 product.

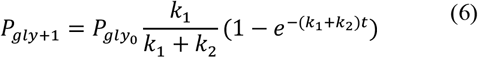

Finally, substituting equation (5) into equation (2) and integrating gives an expression for the consumption of native RNA primer.

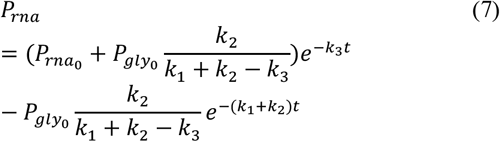

Given independent estimates for *P_gly_0__, k_2_*, and *k*_3_, the rate constant *k*_1_ was estimated from normalized, integrated gel band intensity for P and P_gly+1_ by nonlinear fitting to the system of closed-form solutions (5), (6), and (7). Propagated error on the estimate for *k*_1_, based on errors on the measurements for *P_gly_0__, k*_2_, and *k*_3_, was simulated by the Monte Carlo method^21^.

With C*C dinucleotide (Figure S7):

Primer extension was quantified for each time point by integrating band intensity in each gel lane. Band intensity was normalized in each lane. Remaining primer (P) at each time point, starting from the initial fraction of primer (P_0_), was plotted as −ln(P/P_0_) vs. reaction time, and the observed rate constant, k_obs_, was estimated by the slope of a linear regression line.

With G*U dinucleotide (Figures 6 and S12):

Performed exactly as in the preceding paragraph.

### Base hydrolysis of the +1 NP product (Figure 3B)

A primer extension reaction performed at 2.5 mM MgCl_2_ was allowed to proceed for 40 minutes before being quenched with 29 μL quench buffer (final quench concentrations: 50 mM EDTA, 2 μM reverse complement of the template, 90 % v/v formamide). 2 μL of 1.5 M NaOH was added to 24 μL of the quenched reaction, which increased the pH of the quenched primer extension reaction to 12. The alkaline reaction was incubated at room temperature for 30 s before being neutralized with 1 μL of 0.5 M HCl to pH 8. 3 μL aliquots were subjected to 20 % polyacrylamide gel electrophoresis (19:1 with 7 M Urea) and imaged as the aforementioned primer extension reactions.

### Chemical N-acetylation of aminoacylated primers (Figure 3C)

4 μL of 180 mM Sulfo-NHS-Acetate was added to a typical aminoacylation reaction performed at a 40 μL scale and the reaction was incubated for 2 hours at room temperature. The reaction was precipitated with 0.1 volumes of 5 M NH_4_OAc and 3 volumes of isopropanol on dry ice for 20 minutes and pelleted at 15,000 rpm for 15 minutes at 4 °C. The pellet was washed twice with 80 % ethanol and resuspended in 21 μL nuclease-free water. 10 μL of the precipitated reaction was subjected to the primer extension procedure described in the preceding paragraphs. Because MgCl_2_ from the aminoacylation reaction was washed away during precipitation, MgCl_2_ was added to a final concentration of 2.5 mM in the final reaction. In addition, exact primer concentrations after precipitation could not be accurately determined due to the presence of dFx Flexizyme. Primer concentrations were estimated based on a mock aminoacylation reaction in which dFx Flexizyme was omitted. The mock aminoacylation reaction was subjected to Sulfo-NHS-Acetate labeling as described above and used as the “RNA” control.

### Biotin modification of primer extension reactions (Figure 3D)

Primer extension reactions were performed as described in the preceding paragraphs. At indicated time points, 1 μL of the reaction was quenched in 14 μL of freshly prepared NHS-Biotin buffer (33 mM EDTA, 1 mM NHS-Biotin final after quenching) and incubated at room temperature for 1 h to allow for biotin labeling. At 1 h, 15 μL of quench buffer (final quench concentrations = 26 mM EDTA, 1 μM reverse complement of the template, 46 % v/v formamide) was added to each labeling reaction. The quenched reactions were heated at 92 °C for 2 minutes and 3 μL aliquots were loaded into 20 % polyacrylamide gels (19:1 with 7 M Urea). The gels were run at 20 W for 1 h 20 minutes, imaged, and quantified as the aforementioned primer extension reactions.

### Ligation reactions (Figures 5 and S8-11)

Aminoacylation reactions that were used in subsequent ligation experiments were performed with dFx Flexizyme mutant M2 (Table S1), which recognizes the 3’ terminal ACA sequences. RNA template was added to a typical 10 μL aminoacylation reaction to the final concentration of 3.75 μM, followed by Na-HEPES pH 8.0 to a final concentration of 200 mM. Note: because aminoacylation reactions were performed in the presence of 10 mM MgCl_2_, they contributed 2.5 mM of MgCl_2_ to the final reaction. The reactions were allowed to warm up to room temperature for 2 minutes before being initiated by the addition of the 2-methylimidazole activated decamer (Ligator1, Table S1) to a final concentration of 10 μM. Final reaction concentrations: 2.5 μM mixture of primers, 2.5 mM MgCl_2_, 200 mM HEPES pH 8.0, and 10 μM activated decamer. Reactions were performed in technical triplicates. At indicated time points, 1 μL of each reaction was quenched with 29 μL quench buffer (final quench buffer concentrations: 50 mM EDTA, 2 μM reverse complement of the template, 90 % v/v formamide). Prior to loading on 20 % polyacrylamide gels (19:1 with 7 M Urea), the quenched reactions were heated at 92 °C for 2 minutes to denature the duplex. 3 μL aliquots were loaded into gels and run at 20 W for 1 h 20 minutes. The gels were imaged on Typhoon 9410 imager and band densities quantified in ImageQuant TL software.

To independently determine the kinetics of pure RNA ligation, ligation reactions were performed with the non-aminoacylated RNA primer. The non-aminoacylated primer was subjected to typical aminoacylation conditions, except dFx Flexizyme M2 was replaced with water for those reactions. The ligation reactions were then set up exactly as in the preceding paragraph. Note: to control for possible effects of the different amino acid-DBE esters in the RNA ligation reactions, RNA control reactions were performed in the presence of each tested amino acid-DBE ester (see Figure S8H).

Hydrolysis rates of aminoacylated primers were measured under the same ligation conditions described above, except that unactivated 10mer was used instead of the 2-methylimidazole activated one. The hydrolysis reaction was quenched using the acidic quench buffer (10 mM EDTA, 100 mM NaOAc pH 5.0, 150 mM HCl, 2 μM reverse complement of the template, 70 % v/v formamide), heated at 92 °C for 2 minutes to denature the duplex, and run on an acidic 20 % polyacrylamide gel (19:1 with 7 M Urea, 0.1 M NaOAc pH 5.0).

### Kinetic analysis of ligation reactions

RNA reaction: primer extension was quantified for each time point by integrating band intensity in each gel lane. Band intensity was normalized in each lane. Remaining primer (P) at each time point, starting from the initial fraction of primer (P_0_), was plotted as −ln(P/P_0_) vs. reaction time, and the observed rate constant, k_obs_, was estimated by the slope of a linear regression line. This k_obs_ corresponds to k_3_ in the kinetic model used to obtain k_obs_ (k_1_) of the aminoacylated primer.

Hydrolysis reaction: hydrolysis was quantified for each time point by integrating band intensity in each gel lane. Band intensity was normalized in each lane. Remaining primer-gly (P) at each time point, starting from the initial fraction of primer-gly (P_0_), was plotted as −ln(P/P_0_) vs. reaction time, and the observed rate constant, k_obs_, was estimated by the slope of a linear regression line. This k_obs_ corresponded to k_2_ in the kinetic model used to obtain k_obs_ (k_1_) of the aminoacylated primer.

Aminoacylated reaction: performed using the model described for the primer extension of aminoacylated primers with the following modifications: because the gel band corresponding to the ligation product of aminoacylated primers could not be resolved from the gel band corresponding to the ligation product of pure RNA primers, only time points at which pure RNA primers produced less than 2 % of the ligated product band were used to model the aminoacylated primer ligation.

### Hydrolysis reactions (Table 1, Figures S3 and S4)

Purified products **1** and **2** were subjected to primer extension conditions in single-stranded and double-stranded states without the addition of activated dinucleotides or decamers. Double-stranded reactions were annealed by heating them to 70 °C for 3 minutes and slowly cooling to 20 °C at 0.1 °C/s (singlestranded reactions were not subjected to annealing). Hydrolysis reaction conditions: oligonucleotide (**1** or **2**) 0.375 μM, Na-HEPES pH 8.0 200 mM, MgCl_2_ 2.5 mM or 100 mM, 22 °C (thermocycler). The reactions were stopped by the addition of the quench buffer (final quench concentrations: 50 mM EDTA, 2 μM reverse complement of the template, 90 % v/v formamide) and flash frozen in liquid nitrogen. The doublestranded quenched reactions were heated for 2 minutes at 92 °C to denature the duplex before loading into the gel (the singlestranded reactions were not heated).

**Table 1.** Hydrolytic stability of terminal (1) or internal (2) 2’(3’)-glycyl, phosphoramidate linkage.

|  | Half life (h<sup>-1</sup>) |  |
| --- | --- | --- |
|  | ssRNA | dsRNA |
| **terminal N-P linkage (1)** |  |  |
| 100 mM MgCl<sub>2</sub> | 10.9(3) | 20.2(3) |
| 2.5 mM MgCl<sub>2</sub> | 24.3(4) | 53(1) |
| **internal N-P linkage (2)** |  |  |
| 100 mM MgCl<sub>2</sub> | 8.6(6) | 22.8(2) |
| 2.5 mM MgCl<sub>2</sub> | 20.1(9) | 72(2) |

Kinetic analysis: hydrolysis was quantified for each time point by integrating band intensity in each gel lane. Band intensity was normalized in each lane. Remaining amino acid bridged oligonucleotide (P; either **1** or **2**) at each time point, starting from the initial fraction of the amino acid bridged oligonucleotide (P_0_; either **1** or **2**), was plotted as −ln(P/P_0_) vs. reaction time, and the observed rate constant, k_obs_, was estimated by the slope of a linear regression line. The half-lives were calculated from the k_obs_ values by the following equation for a first-order process: t_1/2_ = ln(2)/k_obs_.

Degradation product characterization of **2**: the reaction was performed for 24 h in a solution that contained 0.375 μM **2**, Na-HEPES pH 8.0 200 mM, and MgCl_2_ 2.5 mM at 22 °C. After 24 h, the reaction was precipitated with 0.1 V 5 M NH_4_OAc and 3 V isopropanol, pelleted at 15,000 rpm at 4 °C, washed twice with 80 % ethanol, desalted using a C18 Zip-tip column, and analyzed on an Agilent 1200 HPLC coupled to an Agilent 6230 TOF mass spectrometer.

## RESULTS

In order to ask whether RNA aminoacylation would interfere with or potentiate RNA copying chemistry, we first established a primer extension assay for RNA copying initiated from a 2’(3’) aminoacylated RNA primer (Figure 2A). To aminoacylate primers we used Flexizyme, a ribozyme originally evolved by *in vitro* selection to aminoacylate any tRNA of interest with a wide variety of amino acids^22,23^. Flexizyme dFx can aminoacylate any RNA sequence that ends in the CA-3’ sequence found in tRNAs^23^. Although aminoacylation occurs at the 3’ hydroxyl, rapid transacylation yields a dynamic mixture of 2’- and 3’-aminoacylated regioisomers^24^. We designed a 10 nt primer terminating in CCA and tested it for aminoacylation using dFx and dinitrobenzyl activated glycine, the dinitrobenzyl group providing the recognition element for this class of Flexizyme. Acylation yields ranging from 26-60 % were obtained, as assessed by polyacrylamide gel electrophoresis (PAGE) under acidic conditions (Figure S1).

To investigate the ability of the 2’(3’)-glycyl oligonucleotide to act as a primer, we designed a template that provides a single binding site for a 5’-5’ aminoimidazolium-bridged cytidine dinucleotide (C*C), the reactive species in primer extension using 2-aminoimidazole activated cytidine ribonucleotides (Figure 2B)^25^. We diluted the RNA acylation reaction mixture into primer extension buffer, added the template strand and MgCl_2_ (2.5 mM) and initiated primer extension by addition of the C*C dinucleotide (20 mM). Note that due to the short halflife of the ester linkage of 2’(3’)-glycyl RNA under primer extension conditions (Figure S5, E and F), and the fact that acylation does not proceed to completion, a mixture of 2’(3’)-glycyl and native 2’,3’-hydroxyl terminated RNA was always present in our Flexizyme-treated reaction mixtures. These two primer species are not separated by PAGE employing Tris-borate EDTA (TBE) buffer, although the products of primer extension can be resolved. Upon analysis of the reaction mixture, two new bands were observed, which we hypothesized to be due to +1 extension from either 2’(3’)-glycyl RNA or native RNA (Figure 2B). Control reactions without either Flexizyme or the glycyl dinitrobenzyl ester displayed only one product band. Further, the inclusion of either Flexizyme or amino acid dinitrobenzyl ester did not interfere with the rate of primer extension at the concentrations employed (*vide infra*).

In principle, primer extension of the aminoacylated primer by reaction with the activated 5’-phosphate of the incoming imidazolium-bridged dinucleotide could occur in either of two ways (Figure 3A). Attack of the free amino group of the 2’(3’)-glycyl RNA would result in the formation of a phosphoramidate linkage, while attack of the remaining 2’(3’)-hydroxyl of the primer would lead to the formation of a phosphodiester linkage. To first confirm the retention of an aminoacyl ester linkage in the reaction products, we subjected the reaction mixture to transient strongly basic conditions (Figure 3B). Treatment with 115 mM NaOH (pH 12) for 30 seconds led to disappearance of the novel (top) +1 band, while leaving the +1 band due to extension from 2’,3’-hydroxyl terminated RNA (bottom) unaffected, indicating the presence of a base-sensitive linkage in the top band. To test whether the free glycyl amino group was needed for extension, we performed primer extension reactions in which the acylated primer was treated with an acetylating reagent (sulfo-NHS-acetate) before the addition of the C*C imidazolium-bridged dinucleotide (Figure 3C). 2.5 mM MgCl_2_ was used in these assays to allow detection of both +1 bands at earlier time points, as the unmodified RNA primer reacts slowly under these conditions. Reaction mixtures thus treated were identical to those obtained from control reactions lacking Flexizyme, indicating that the nucleophilic glycyl amino group is required for formation of the novel +1 product. Finally, to directly prove that the free amino group is consumed during primer extension, we employed biotin labeling (Figure 3D). In this experiment, NHS-biotin was added to the primer extension reaction quench solution at each time-point. If a free amino group is present, NHS-biotin will react, leading to a clear gel shift. Conversely, if no amino group is present due to N-P bond formation, no gel shift due to biotin labeling is possible. As seen in Figure 3D, as the reaction with the 2’(3’)-glycyl RNA proceeds, the extent of labeling with biotin decreases as the +1 band increases in intensity (left panel). No labelling is observed with chemically acetylated 2’(3’)-glycyl RNA (right panel) or 2’,3 ‘-hydroxyl terminated RNA (middle panel). Taken together, the results from base hydrolysis and chemical labeling experiments strongly support the formation of a phosphoramidate linkage during primer extension from 2’(3’) glycyl RNA.

To investigate the reactivity of 2’(3’)-aminoacylated, phosphoramidate-linked RNA (‘amino acid bridged RNA’), we required a means of isolating single-stranded RNA containing site-specific phosphoramidate linkages, to study primer extension, ligation, and hydrolysis. We adapted our recently reported strategy for the generation of site-specific 3’-5’ pyrophosphate linkages to provide a means to access these unusual amino-acid bridged RNAs (Figure S2)^26^. To obtain 2’(3’)-aminoacylated, phosphoramidate-linked RNA containing only a single-nucleotide extension (‘terminal’ amino acid bridged RNA), we first aminoacylated an RNA primer using Flexizyme dFx. Following acylation, we performed a primer extension reaction using a DNA:RNA hybrid template in which the region to be copied is RNA but the remainder of the template is DNA. The template contains only a single binding site for the activated C*C imidazolium-bridged dinucleotide. Incubating the primer template duplex with C*C dinucleotide leads to robust conversion of the primer to extended products in which the +1 nucleotide is connected to the terminal amino group of the 2’(3’)-aminoacylated RNA by a phosphoramidate linkage. We omitted Mg^2+^ from the primer extension reaction to discourage extension of the fraction of the RNA primer that was not acylated in the Flexizyme reaction. DNase digestion of the template then facilitated recovery of the modified primer, which we purified using preparative gel electrophoresis. To obtain RNA strands in which the amino acid linkage is followed by a longer stretch of ribonucleotides, we replaced the primer extension reaction with a ligation reaction, using a ligator RNA oligonucleotide bearing a 2-methylimidazole group activating the 5’-phosphate. In this case, the entire template was DNA so that following ligation, DNase treatment and preparative gel electrophoresis enabled purification of the modified, amino acid linked strand.

We compared the chemical stability of amino acid bridged RNA to that of canonical RNA, to determine whether it could, in principle, support the replication of genetic information and enable ribozyme function. Liu et al. previously measured a pH-rate profile for the breakdown of a methyl-tyrosine linked dinucleotide as a model for the behavior of amino-acid bridged RNA co-polymers^9^. The reported optimum stability at pH ~6.0 reflects a balance between acid-catalyzed hydrolysis of the phosphoramidate bond and base-catalyzed hydrolysis of the aminoacyl ester linkage. The stability of the aminoacyl ester linkage was greatly enhanced upon phosphoramidate formation, potentially providing a mechanism for stable capture of amino acids by RNA strands in a prebiotic setting.

Next, we examined the stability of longer amino acid bridged RNAs under conditions relevant to RNA copying chemistry. We first prepared two single-stranded RNAs containing either a terminal (AGAGAAGCAA-gly-C, **1**) or internal 2’(3’)-glycyl phosphoramidate linkage (AGAGAAGAGAGCAGACA-gly-CCCGGCAGCU, **2**), using the strategies outlined above. The 2’(3’)-glycyl, phosphoramidate-linked RNAs were then incubated at 22 °C in a pH 8.0 solution (conditions typical for primer extension reactions) at either high (100 mM) or low (2.5 mM) concentrations of Mg^2+^, and in the presence or absence of a complementary strand (Table 1, Figure S3). Maximum stability was observed for duplex products at low concentrations of Mg^2+^. Under these conditions, we observed half-lives of 53 hours for terminal linkages and 72 hours for internal linkages. In the presence of 100 mM Mg^2+^, conditions employed for template copying experiments (see below) the stability was reduced, although the observed half-lives on the order of hours are still sufficient to enable amino acid linked RNAs to act as templates for further copying cycles (*vide infra*). All reactions were performed at pH 8.0, 200 mM HEPES. Values are reported as the mean +/- standard deviation from triplicate experiments.

As the gel-based assay does not report on the specific linkage cleaved (ester vs. phosphoramidate), we desalted samples from the cleavage of **2** after 24 hours and analyzed the reaction products by LC-MS. Only products resulting from ester cleavage could be observed (Figure S4), consistent with the previously reported stability of the phosphoramidate linkage at pH values above 5.0.^9^

The proportion of amino acid bridged RNAs within a prebiotic population of polynucleotides would depend on the rates of amino acid activation and aminoacylation of RNA, and the competing rates of aminoacyl ester hydrolysis and phosphoramidate linkage formation. To investigate the rate of primer extension via phosphoramidate formation we measured the rates of primer extension on a template designed to provide a single binding site for C*C imidazolium-bridged dinucleotide (Figure 4, Figure S5). To quantify the kinetics of glycyl-RNA primer extension, we used a simplified model of the reaction (for details, see Supporting Information). At the start of the reaction, due to incomplete acylation by Flexizyme, both an aminoacylated and non-acylated primer species are present, which are not resolvable by gel electrophoresis. We monitored the primer extension reaction by PAGE, which can resolve the two different +1 extended reaction products, with or without a bridging amino acid residue. To follow the reaction kinetics, we modeled the hydrolysis of the aminoacylated primer to the native RNA primer as a first-order irreversible reaction with a rate constant k_2_. The pseudo first-order rate constant, k_1_, for extension of the aminoacylated primer was estimated by nonlinear regression using independently measured values for the aminoacyl hydrolysis rate constant, k_2_, the native RNA primer extension rate constant, k_3_, and the initial fraction of aminoacylated primer Pgly_0_ (Figure S5).

By following the procedures outlined above, we obtained estimated rates for primer extension reactions using primers terminated in either a 2’-3’ cis diol or a 2’(3’)-glycyl group (Figure 4, Figure S5). We note that the rates we report combine possible reactions initiated from both 2’- and 3’-linked glycyl residues. At pH 8.0 and 50 mM Mg^2+^ the rate of primer extension via phosphoramidate bond formation was similar to that observed for phosphodiester bond formation (k_1_ = 11 h^-1^ vs. 9 h^-1^ for RNA). We have previously observed that N-P bond formation using 2’ or 3’ amino groups is insensitive to Mg^2+^ concentration^12,13^. This feature is highly desirable if genetic copying chemistry is to be integrated within fatty acid vesicles, as concentrations of free Mg^2+^ above 4 mM degrade and precipitate such membranes. We therefore performed the same primer extension reactions with and without 2.5 mM Mg^2+^. The observed rates for primer extension via phosphoramidate bond formation were insensitive to Mg^2+^, whereas phosphodiester bond formation was much slower at lower Mg^2+^ concentrations. In the absence of Mg^2+^, the rate of primer extension for the 2’(3’)-glycyl terminated primer was 2 orders of magnitude greater than for extension from the canonical diol-terminated RNA (k_1_ = 12.0 h^-1^ vs. 0.118 h^-1^ for RNA). These results are in accordance with the greater nucleophilicity of the amino substituent relative to the hydroxyl group and the presumed requirement for divalent metal mediated deprotonation of the 3’-hydroxyl to afford the Mg-bound alkoxide, the most likely active species for primer extension of canonical RNA. The much greater reactivity of the glycyl-terminated RNA raised the question of whether primer extension from this modified primer is still template dependent. However, we found that primer extension in the presence of the template yields 76% extended product after 15 minutes compared to only 6% after 30 minutes in the absence of the template (Figure S6).

The incorporation of mismatched bases^27^ and non-canonical nucleotides^28^ can stall primer extension, presumably due to suboptimal geometry of the reaction center. To see if an amino acid bridge would interfere with proper pairing of the terminal primer-template base pair, we quantified the rate of reaction for a primer in which the terminal 3’ nucleotide is joined by a phosphoramidate linkage to an upstream glycine ‘bridge’ (Figure S7). The observed rate for extension downstream of the amino acid bridge was similar to that obtained for an identical primer containing only phosphodiester linkages (10.7 h^-1^ vs 8.8 h^-1^ for RNA). Thus, the incorporation of a bridging amino acid does not significantly retard downstream primer extension steps.

In addition to the polymerization of activated nucleotides, RNA templates can also be copied by the ligation of short oligomers^16^. This scenario is attractive as ligation requires fewer chemical reaction steps than primer extension to copy a template of given length. However, rates of RNA ligation are much lower than for polymerization^19^; consequently, loss of the activating group competes with ligation, leading to overall low yields. The slow rate of RNA ligation can be explained by the fact that the leaving group is simply a protonated imidazole, which lacks the highly preorganized structure of the imidazolium-bridged dinucleotide intermediate of primer extension. In fact, short oligoribonucleotides ending with 3’-amino-2’,3’-dideoxy-ribonucleotides show ligation rates that are orders of magnitude faster than all-RNA oligonucleotides^16^. However, no potentially prebiotic route to 3’-amino nucleotides is yet known. We therefore wondered whether the more prebiotically plausible 2’(3’)-aminoacylated RNA would show similar trends in rate and yield for non-enzymatic ligation.

To compare the rates of ligation of unmodified RNA and aminoacylated RNA, we used a ligation assay similar to that developed for primer extension and we used the same formalism to model the kinetic system, assuming saturation of the primer-template duplex with ligator. For ligation reactions, we were able to determine the rate of formation of ligated product from aminoacylated RNA by collecting data at time-points where reaction with the control RNA primer was negligible. As input to our kinetic model we measured the rates of hydrolysis and the rates of control reactions with an RNA primer in the presence of the amino acid dinitrobenzyl ester (Figure S8).

We first tested the template-directed ligation of a 2’(3’)-glycyl primer with either a 2-methylimidazole or 2-aminoimidazole activated decamer (Figure S9). Although 2-aminoimidazole is a superior activating group for non-enzymatic polymerization, due to enhanced formation of the active imidazolium bridged dinucleotide intermediate, 2-methylimidazole activation is superior for N-P ligation^16^. This is consistent with the fact that 2-aminoimidazole is an intrinsically worse leaving group, due to its higher *pK_a_*^29^. Indeed, in our system, the observed rate of ligation was 6-fold greater for the 2MeI activated ligator (1.81 h^-1^ vs 0.281 h^-1^). Notably, the rate of ligation for the 2’(3’)-glycyl primer reacting with a 2-methylimidazole activated ligator, at 2.5 mM Mg^2+^, was ~500 times greater than for the equivalent reaction with unmodified RNA (Figures 5 and S9). In the absence of template, no ligation was observed on the time-scale of the experiment (Figure S10).

**Figure 5.**
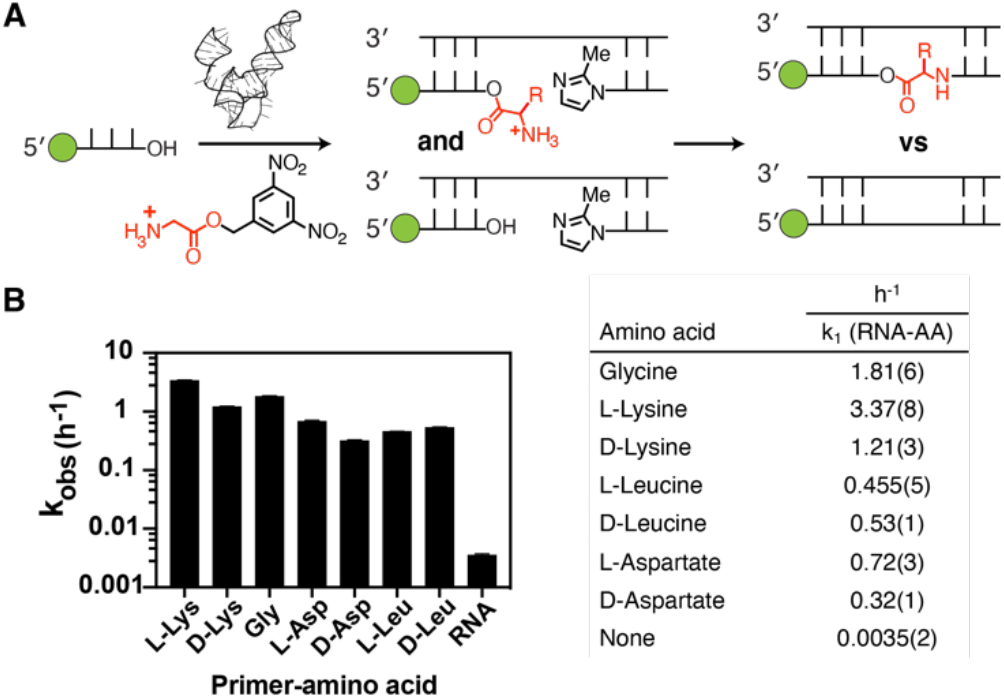
Kinetic analysis of nonenzymatic ligation from 3’ hydroxyl and 2’(3’) aminoacyl primers. **(A)** Schematic representation of the assay. **(B)** Kinetic parameters. The k_obs_ (h^-1^) values are plotted against the primer-amino acid tested. ‘None’ refers to the unmodified RNA reaction in the presence of DBE-glycine but the absence of Flexizyme. The pseudo first-order rate constant, k_1_, for ligation of the aminoacylated primer was estimated by nonlinear regression using values for k_3_, the native RNA ligation rate constant, k_2_, the aminoacyl hydrolysis rate constant, and the initial fraction of aminoacylated primer Pgly0, that were measured independently. Plots used to obtain k_1_, k_2_ and k_3_ can be found in Figure S8. Uncertainty in the estimate of k_1_ was analyzed by the Monte Carlo method to propagate error on the estimates of independent model parameters. All reactions were performed at pH 8.0, 200 mM HEPES, 2.5 mM MgCl_2_. Values are reported as the mean +/- standard deviation from triplicate experiments.

Having demonstrated enhanced ligation rates with aminoacylated RNA, we were interested in determining whether the ligation rate would differ significantly across a panel of amino acids (Figure 5, Figure S8). We tested 8 amino acids that differ in charge, size and stereochemistry. All amino acids, except for *N*α-acetyl-L-lysine, reacted orders of magnitude faster than RNA under the conditions of the assay (pH 8.0 and 2.5 mM Mg^2+^). L-Lys reacted at the greatest rate, which is perhaps surprising given its length and positively charged side chain. Acetylation of the lysine α-amino group blocked ligation completely while acetylation of the ε-amino group reduced the rate 3-fold, confirming regioselectivity for reaction of the α-amine versus the ε-amine (Figures S8 and S11). Lysine and aspartate both displayed a preference for reaction of the L-enantiomer, although leucine displayed no such preference. Overall, it is notable that amino acids with different properties could be incorporated into RNA via ligation. For example, the carboxyl and amino side chains introduced by aspartate and lysine, respectively, have no parallel in native RNA. Such integration of novel functionality may allow for the expansion of the catalytic repertoire of ribozymes assembled by non-enzymatic ligation.

To determine whether an RNA strand containing a single amino acid bridge could act as a template for RNA primer extension we used the bridged 27mer ssRNA **2** (5’-AGAGAA-GAGAGCAGACA-gly-CCCGGCAGCU-3’), and an all-RNA control, as templates. The bridged template contains a stretch of residues 5’-ACA-gly-C-3’ such that only one activated dinucleotide intermediate, G*U, was necessary to compare template directed copying at three positions. We tested three cases that differed only in the position of the primer 3’-end relative to the glycine bridge in the template. In the first case, the imidazolium-bridged G*U dinucleotide spans the glycine bridge (Figure 6A). In the second case, the glycine bridge is located after the primer annealing site (Figure 6B). In the third case, the primer extends over the glycine bridge (Figure 6C). Using 20 mM activated G*U dinucleotide and 100 mM Mg^2+^, we evaluated copying in all three cases. We observed primer extension in all three cases, implying that glycine-bridged RNA can indeed act as a template for RNA copying. We estimate that the kinetic defect due to copying over glycine-bridged RNA is approximately 4-fold relative to an all RNA control template for all three cases (Figure 6A-C, Figure S12).

**Figure 6.**
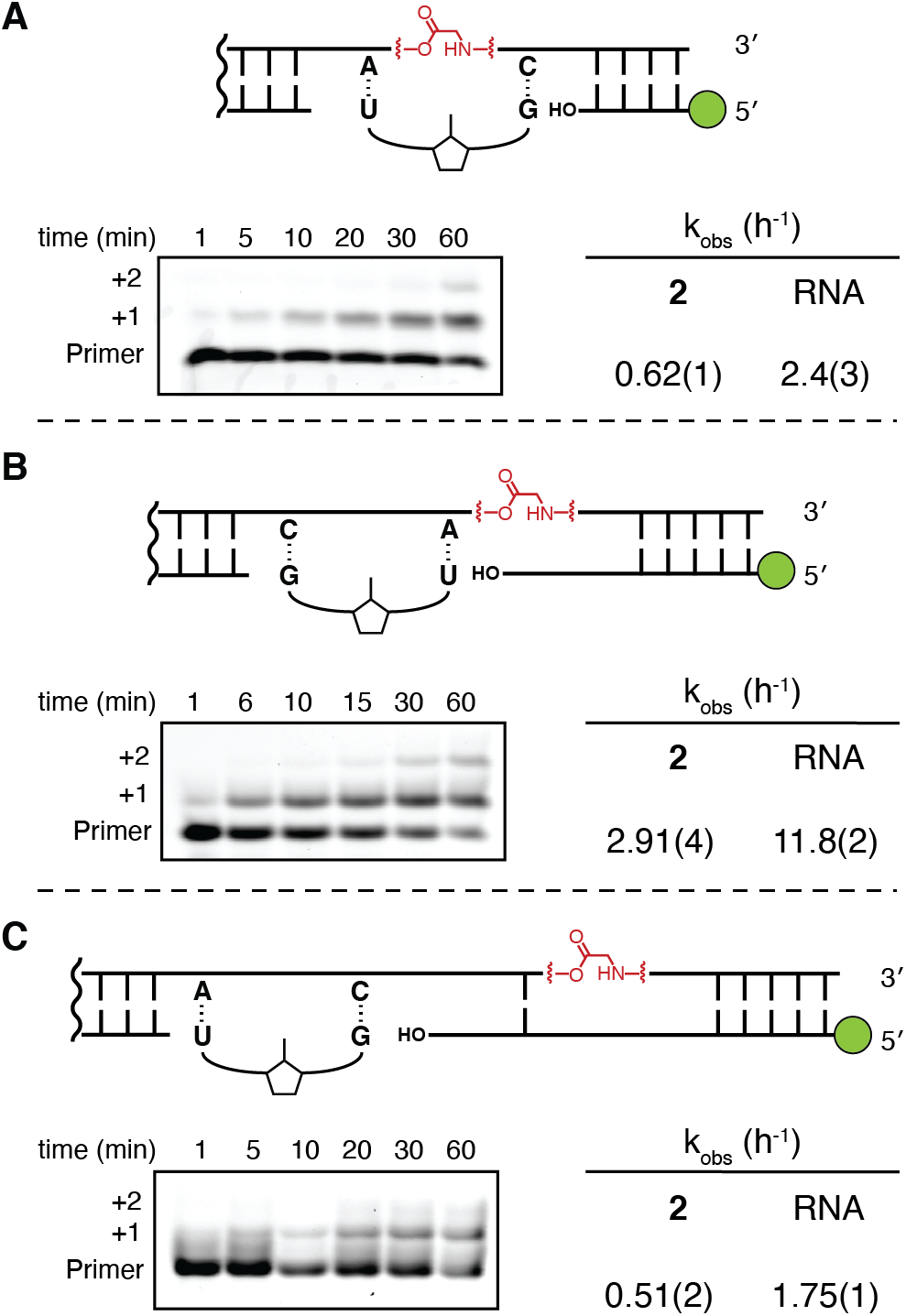
**(A-C)** Kinetic analysis of nonenzymatic primer extension across an RNA template containing a single glycine amino acid bridge. Top: Schematic representation of the primer-template duplexes analyzed, showing the binding site for the G*U dimer in each case. RNA controls contained the identical primer, with the template identical except for the absence of the bridging amino acid. Bottom: Time course of primer extension as monitored by polyacrylamide gel electrophoresis. **2** = ssRNA template with a single glycine bridge. All reactions were performed at pH 8.0, 200 mM HEPES, 100 mM MgCl_2_, with 20 mM G*U dimer. Gel images of RNA control reactions and plots used to determine kinetic parameters are in the Supplementary Information Figure S12.

## DISCUSSION

We have found that the aminoacylation of an RNA primer can, under certain conditions, greatly enhance the nonenzymatic copying of an RNA template. The reaction of aminoacylated RNA primers with incoming imidazolium-bridged dinucleotides gives RNA products containing an amino acid ‘bridge’ composed of a 5’ (C-terminal) ester linkage and a 3’ (N-terminal) phosphoramidate linkage. Notably, this copying reaction proceeds in the absence of Mg^2+^, which is damaging to protocell membranes at low millimolar concentrations. We have also examined the nonenzymatic, template-directed ligation of an aminoacylated RNA strand to a 2-methylimidizaole activated ligator RNA. In this case, rates of ligation are enhanced by at least two orders of magnitude. Similar rate enhancements are seen with primers terminating in 3’-amino-2’,3’-dideoxy-ribonucleotides^16^, however, no pre-biotic synthesis of 3’-amino nucleotides has been described. In contrast, the aminoacylation of RNA is central to biology.

We employed Flexizyme-catalyzed acylation to obtain high yields of aminoacylated RNA primers for our studies. It is possible that RNA aminoacylation only began to play a significant role in the RNA World after the evolution of ribozymes became widespread, but an earlier process would open up the possibility of a role for aminoacylation chemistry in nonenzymatic RNA replication or ligation-mediated ribozyme assembly. The aminoacylation of RNA has been reported from mixtures of phosphorimidazolide activated nucleotides, imidazole, and amino acids^2^, but these reactions are quite inefficient. The discovery of more effective, prebiotically plausible chemistry for RNA aminoacylation would suggest the potential for a common role for this chemistry in the origins of both replication and translation. An efficient chemical aminoacylation process would also be experimentally useful if it overcame the sequence limitations enforced by our use of the Flexizyme ribozyme, which only acylates RNA terminating in CA-3′.

The regioselectivity of the phosphoramidate forming primer extension and ligation reactions remains unknown. Our gel-based analysis cannot distinguish between reactions initiated from the 3′ and the 2′ esters, because of the rapid transacylation of the initially formed aminoacyl ester. The different regioisomers, if a mixture indeed results from phosphoramidate formation, may display different templating activities and stabilities that have been conflated in this study.

It has been noted previously that the enhanced lifetime of the aminoacyl ester linkage to RNA upon formation of a neighboring phosphoramidate linkage may provide a mechanism for the stable integration of amino acid functionality into RNA^9^. Our stability studies revealed the protective effect of duplex formation, which enhances the kinetic stability of amino acid bridged RNA approximately two-fold. Notably, high concentrations of Mg^2+^ promote degradation of the amino acid ‘bridge’; taken together with the much enhanced rates of RNA copying observed at low concentrations of Mg^2+^, this result suggests that amino acid bridged RNA would accumulate preferentially under conditions of low free Mg^2+^, conditions that are also most favorable for protocell stability. The major products of degradation of an amino acid ‘bridge’ under RNA copying conditions are a 5’ fragment composed of native RNA, resulting from aminoacyl ester cleavage, and a 3’ fragment bearing an amino acid at the 5’ terminus linked by a phosphoramidate linkage. Such 5’-N-linked amino acids have been shown to be highly competent for further extension into peptides under activating conditions^30^. Thus, phosphoramidate bond formation via either non-enzymatic primer extension or ligation, followed by hydrolysis of the aminoacyl ester, could initiate peptide synthesis, in addition to the functions outlined above.

We have shown that RNA containing an amino acid bridge remains competent as a template for further cycles of copying. It remains unknown whether amino acid bridged RNA may serve a catalytic function. Ribozyme function can be enhanced using free amino acids as cofactors^31^. Further, introducing novel functional groups to RNA via chemical modification has proven a powerful approach to obtaining ribozymes with enhanced^32^ or new-to-nature functions^33^. Our results show that non-enzymatic ligation with different amino acids can furnish RNA strands with bridging amino acids with a range of side chains. This novel route to the integration of amino acids within RNA may provide new opportunities for ribozyme catalysis that would be exciting to test.

## ASSOCIATED CONTENT

### Supporting Information

Materials and Methods, Figures S1-S12, Table S1, and Characterization of G*U dinucleotide and 3,5-dinitrobenzyl esters of amino acids. This material is available free of charge via the Internet at http://pubs.acs.org.

## Author Contributions

## Notes

The authors declare no competing financial interests.

## ACKNOWLEDGMENT

J.W.S is an investigator of the Howard Hughes Medical Institute. The authors would like to thank Prof. Yamuna Krishnan and Dr. Saurja Dasgupta for helpful comments on the manuscript, Drs. Harry R.M. Aitken, Lijun Zhou, and Li Li for helpful discussions, and Dian Ding for help with G*U synthesis. This work was supported in part by a grant (290363) from the Simons Foundation to J.W.S., a grant from the NSF (CHE-1607034) to J.W.S., and a grant from NASA (NNX15AL18G) to J.W.S.

