## Supporting Information for "A Potential Role for Aminoacylation in Primordial RNA Copying Chemistry"

**Table of contents**

Supplementary Figures	3

Supplementary Tables	22

Characterization of G*U dinucleotide and 3,5-dinitrobenzyl esters of amino acids	24

**Supplementary Figures**

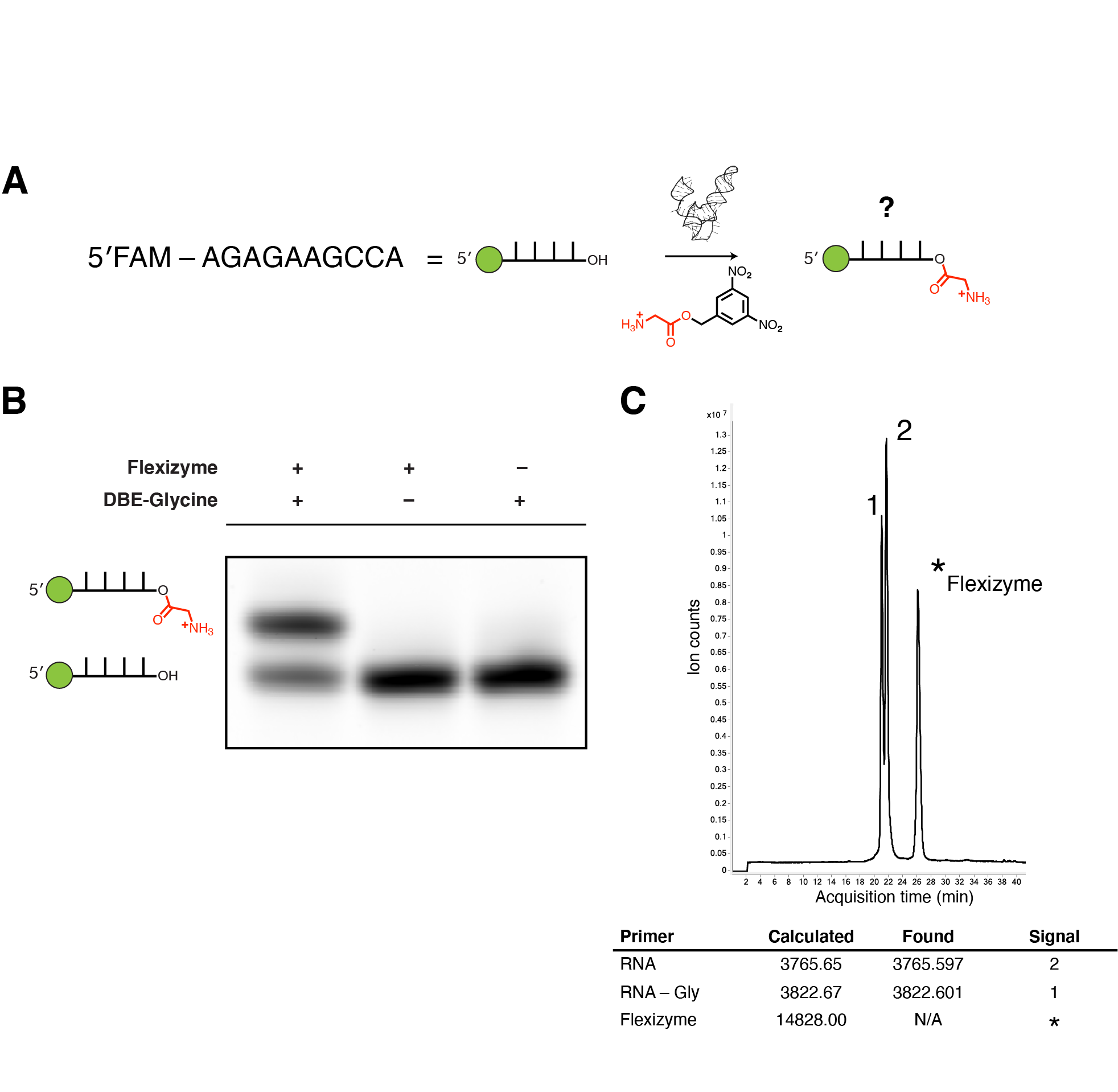

**Figure S1**

Flexizyme-catalyzed aminoacylation of RNA oligomers.

A. Schematic of the strategy used to aminoacylate RNA oligomers. Fluorescently-labeled RNA 10mer is subjected to aminoacylation by the dFx Flexizyme. Amino acid substrate of the dFx Flexizyme is the dinitrobenzyl ester of glycine (DBE-Gly).
B. Aminoacylation was monitored using 20% denaturing acid Urea-PAGE (0.1 M sodium acetate pH 5.0, 7 M Urea; running buffer 0.1 M sodium acetate pH 5.0). Under acidic conditions, the aminoacyl ester bond is sufficiently stable to enable gel resolution of the bands corresponding to the non-aminoacylated and aminoacylated RNA 10mer. Reaction conditions: 50 mM HEPES pH 8.0, 10 mM MgCl_2_, 10 µM RNA 10mer, 5 mM DBE-Gly (20% final DMSO), 10 µM dFx, 14 h incubation on ice.
C. Aminoacylation was monitored using liquid chromatography coupled to a TOF mass spectrometer. Chromatographic resolution yielded 3 distinct signals in the total ion chromatogram. The dominant molecular ions in signals labeled as 1 and 2 were observed to correspond to calculated exact masses of the aminoacylated RNA 10mer and the RNA 10mer, respectively. Signal * did not yield an interpretable m/z, likely due to it being out of the range of the mass detector and is presumed to be dFx.

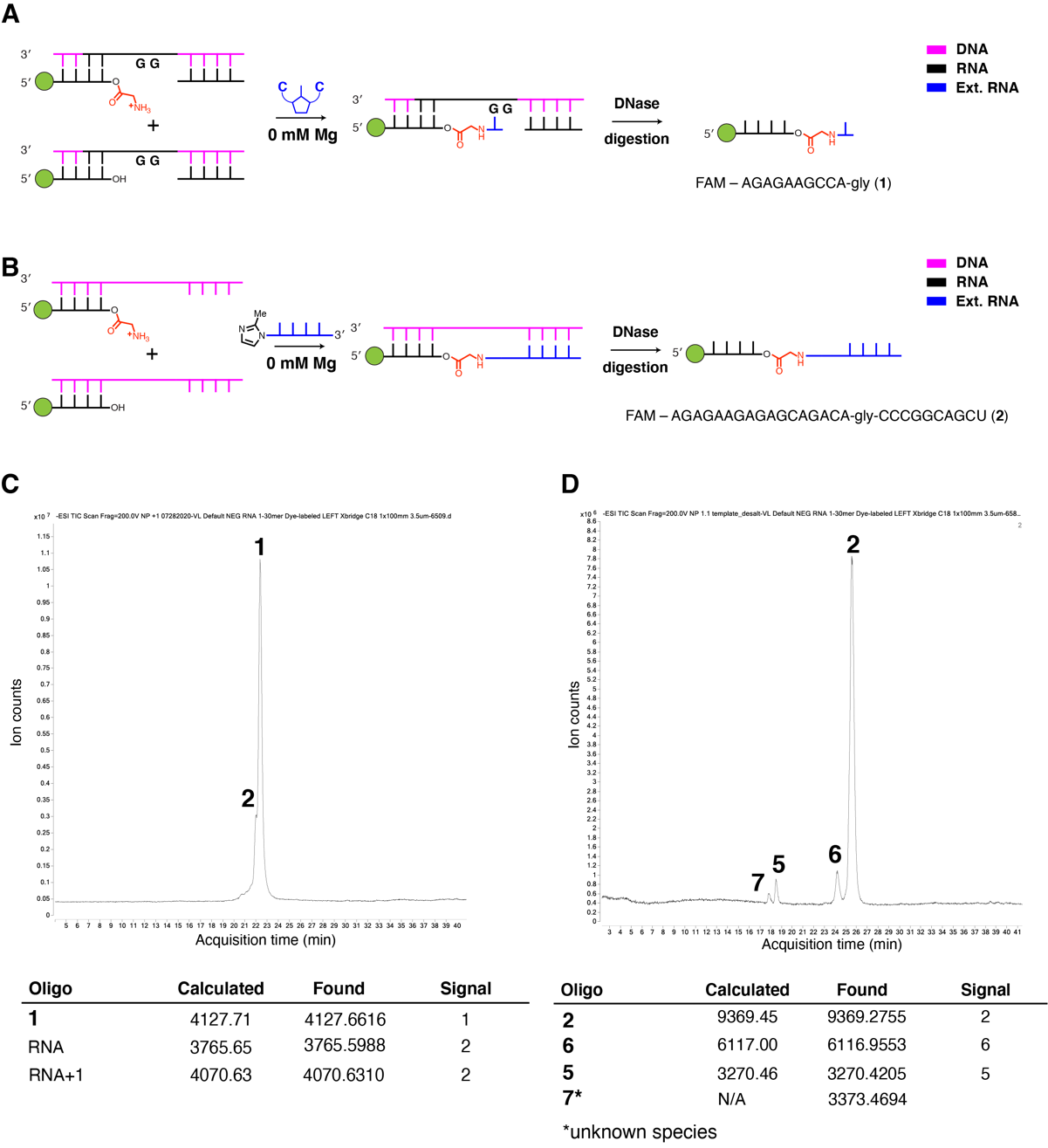

**Figure S2**

Strategy to access glycine-linked +1 and +10 products.

A. Glycine-linked +1 product was accessed by performing a primer extension reaction (200 mM HEPES pH 8.0, with 13.5 mM C*C dimer; trace Mg^2+^ was chelated with a 20-fold excess EDTA to prevent the RNA reaction) on a mixture of aminoacylated and non-aminoacylated fluorescently-labeled RNA 10mer. The template was composed of DNA nucleotides, except for the 4 ribonucleotides in and adjacent to the templating region. TurboDNase was used to digest the DNA template and denaturing 20% Urea-PAGE was used to obtain product **1** with purity of 80-90%.
B. Glycine-linked +10 product was accessed by performing a ligation reaction (standard conditions except Mg^2+^ was chelated with a 20-fold excess EDTA to prevent the RNA reaction) on a mixture of aminoacylated and non-aminoacylated fluorescently-labeled RNA 17mer. The template was fully composed of DNA nucleotides. TurboDNase was used to digest the DNA template and denaturing 20% Urea-PAGE was used to obtain the product **2** with 90% purity.
C. Final product of the reaction shown in A was identified using liquid chromatography coupled to a TOF mass spectrometer. The total ion chromatogram showed two signals that were not fully chromatographically resolved, labeled 1 and 2. The molecular ion identified in signal 1 corresponded to the calculated exact mass of **1**, while the molecular ions in signal 2 corresponded to the calculated exact masses of the starting RNA 10mer and its +1 extension product. Purity of **1** was assessed based on normalized band intensities after gel electrophoresis (data not shown).
D. Final product of the reaction shown in B was identified using liquid chromatography coupled to a TOF mass spectrometer. The total ion chromatogram showed four signals that were fully chromatographically resolved, labeled 2, 6, 5, and 7 (labeling scheme corresponds to the one in Figure S4A). The molecular ion identified in signal 2 corresponded to the calculated exact mass of **2**. The observed molecular ions in signals 6, 5, and 7 and their identities are listed in the table below the chromatogram. Purity of **2** was assessed based on normalized band intensities after gel electrophoresis (data not shown).

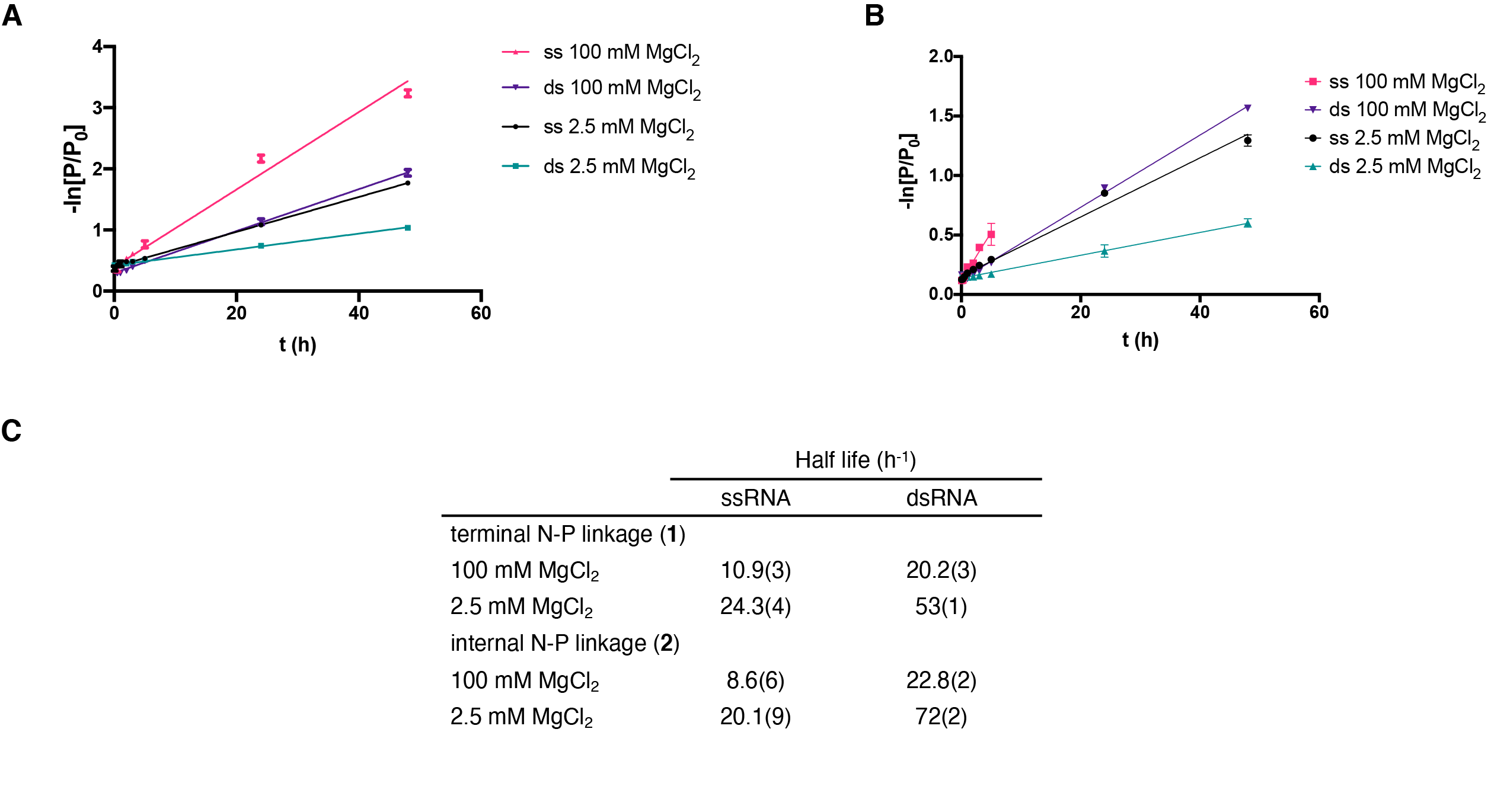

**Figure S3**

Hydrolysis of glycine-linked +1 (**1**) and +10 (**2**) products at different MgCl_2_ concentrations.

A. Hydrolytic stability of the terminal 2′(3′) glycyl phosphoramidate linkage in the glycine-linked +1 product. Shown is the plot of the negative natural logarithm of the ratio of oligomer **1** (P) to initial oligomer **1** (P_0_) versus the time of reaction. The slope of the linear fits corresponded to k_obs_ for each of the tested conditions. Abbreviation ss = single-stranded; ds = double-stranded.
B. Hydrolytic stability of the internal 2′(3′) glycyl phosphoramidate linkage in the glycine-linked +10 product. Shown is the plot of the negative natural logarithm of the ratio of oligomer **2** (P) to initial oligomer **2** (P_0_) versus the time of reaction. The slope of the linear fits corresponded to k_obs_ for each of the tested conditions. Abbreviation ss = single-stranded; ds = double-stranded. The single-stranded reaction at 100 mM MgCl_2_ occurred rapidly enough to permit linear fitting and k_obs_ determination after 5 hours.
C. Summary of the obtained kinetic parameters from A and B. The half-lives were calculated from k_obs_ values by the following equation for a first-order process: t_1/2_ = ln(2)/k_obs_.

All reactions were performed at 200 mM HEPES pH 8.0.

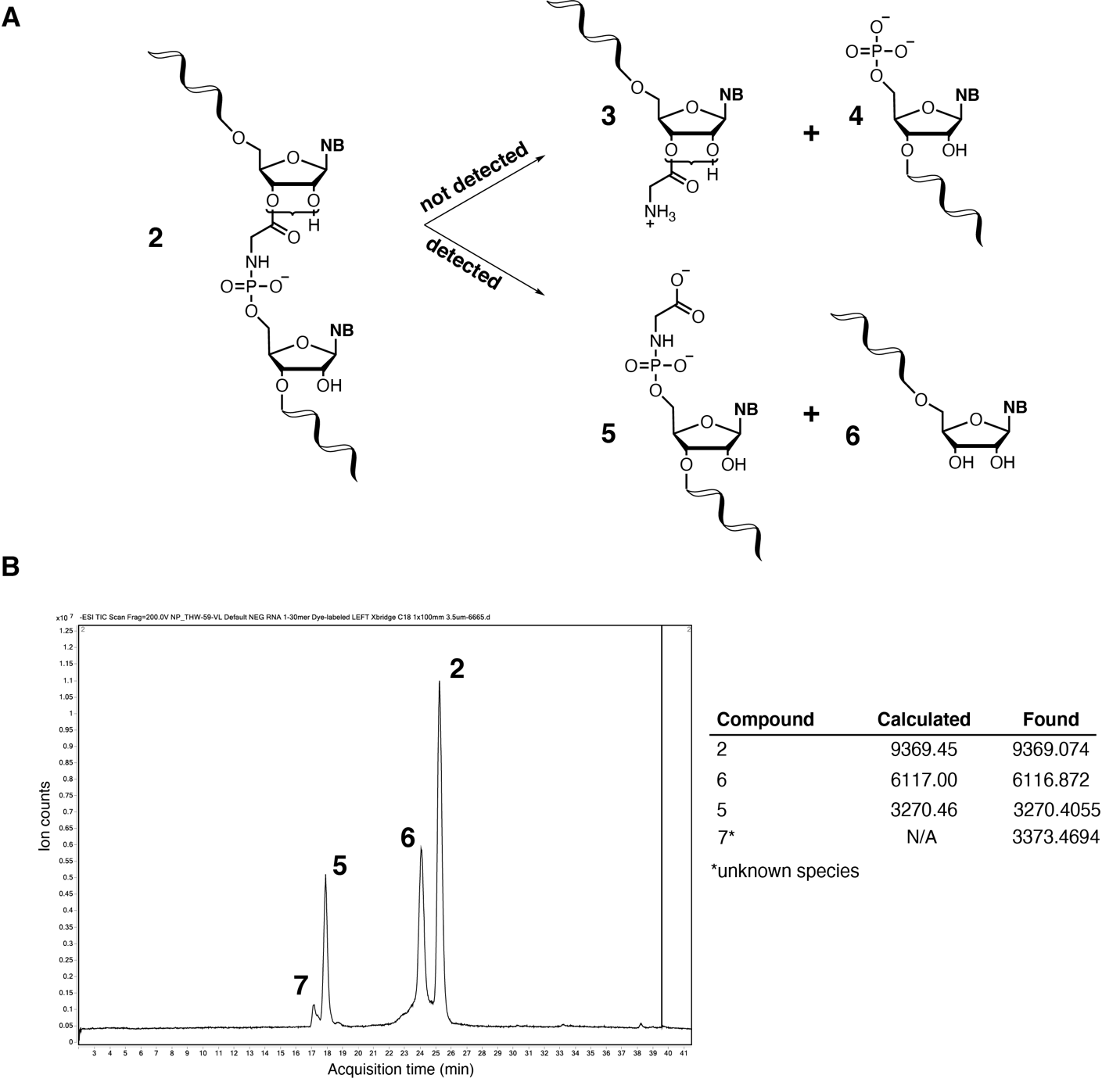

**Figure S4**

Preferential aminoacyl ester linkage hydrolysis of a glycine-linked RNA 27mer (**2**).

A. Potential degradation products of a glycine-linked RNA oligomer. Degradation can occur via the hydrolysis of the aminoacyl ester linkage or the phosphoramidate linkage to yield products that can be distinguished by liquid chromatography coupled to mass spectrometry.
B. Glycine-linked RNA 27mer (**2**) was incubated at pH 8.0, 200 mM HEPES, 2.5 mM MgCl_2_ at 22 ºC for 24 h. The reaction was subjected to liquid chromatography coupled to a TOF mass spectrometer. The total ion chromatogram yielded four distinct, fully chromatographically resolved signals. The dominant molecular ions observed in each signal and their identities based on calculated exact masses are listed in the shown table. The numbering in the chromatogram and the table follows the numbering from A. The observed signals corresponded to the expected products of the aminoacyl ester linkage hydrolysis. No signals that corresponded to the expected products of the phosphoramidate linkage hydrolysis were observed, suggesting that the phosphoramidate linkage is stable under typical primer extension and ligation reaction conditions.

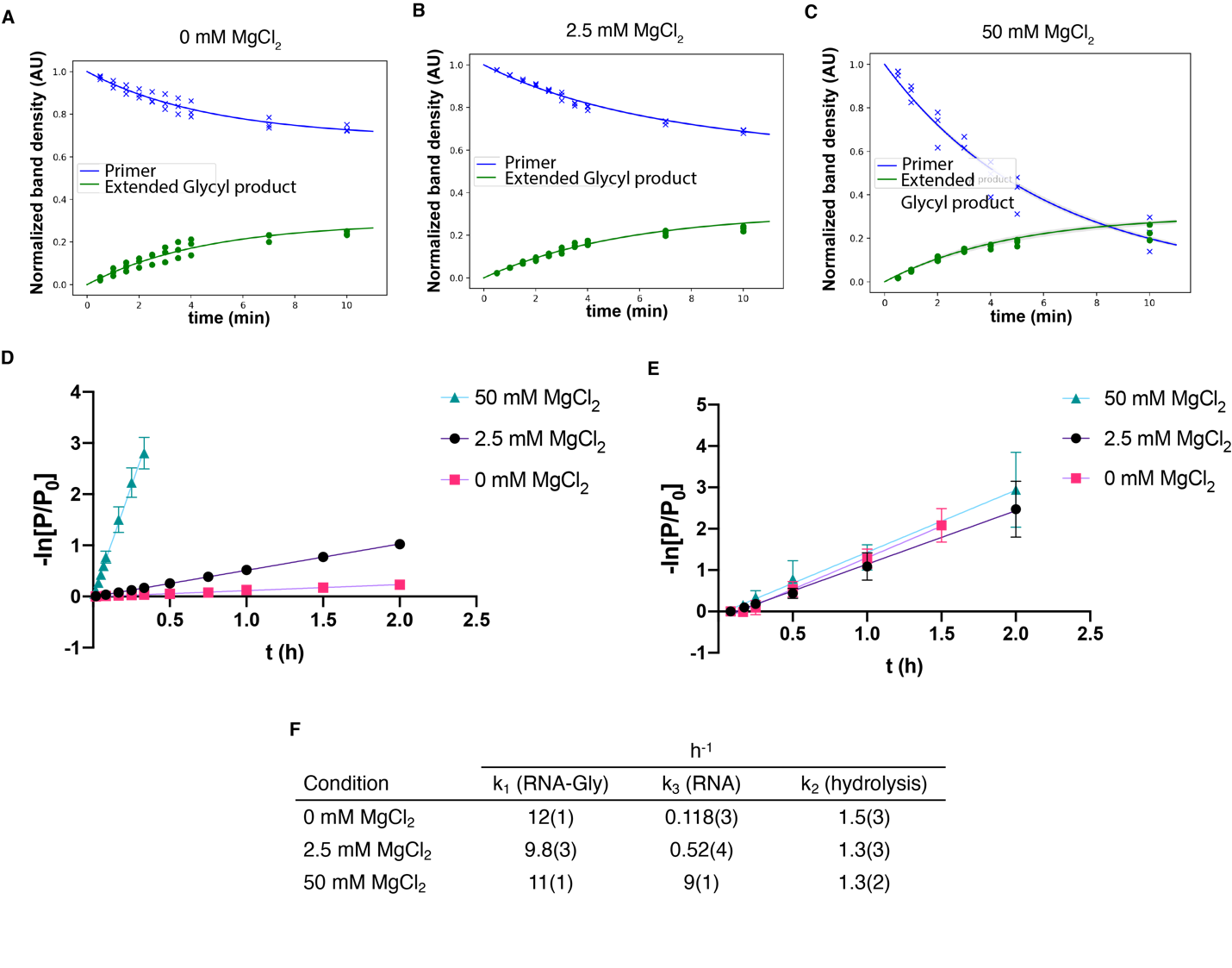

**Figure S5**

Kinetic analysis of nonenzymatic primer extension from 3′ hydroxyl a­­nd 2′(3′) glycyl terminated primers.

A.-C. Plots showing the normalized band intensities for the primer band, which includes both the primer-gly and primer species, (blue) and the extended glycyl +1 product (green) over the course of a primer extension reaction. Non-linear fits were obtained using the kinetic model described in the Materials and Methods section.
D. RNA primer control primer extension reaction. Shown is the plot of the negative natural logarithm of the ratio of primer (P) to initial primer (P_0_) versus time of primer extension for a pure RNA primer at different MgCl_2_ concentrations. The slope of the linear fits corresponded to k_obs_ for RNA reactions that we defined as k_3_ in the kinetic model.
E. Primer-gly hydrolysis reaction. Shown is the plot of the negative natural logarithm of the ratio of primer-gly (P) to initial primer-gly (P_0_) versus time of reaction for the glycine aminoacylated primer at different MgCl_2_ concentrations. The slope of the linear fits corresponded to k_obs_ for primer-gly hydrolysis that we defined as k_2_ in the kinetic model.
F. The tabulated summary of the obtained kinetic parameters from A-E plots. The values are reported as the mean +/− standard deviation from triplicate experiments.

The reactions in A-D were performed at 200 mM HEPES pH 8.0, with 20 mM C*C dimer. The hydrolysis reaction in E was performed at 200 mM HEPES pH 8.0, with the addition of 20 mM CMP instead of the C*C dinucleotide.

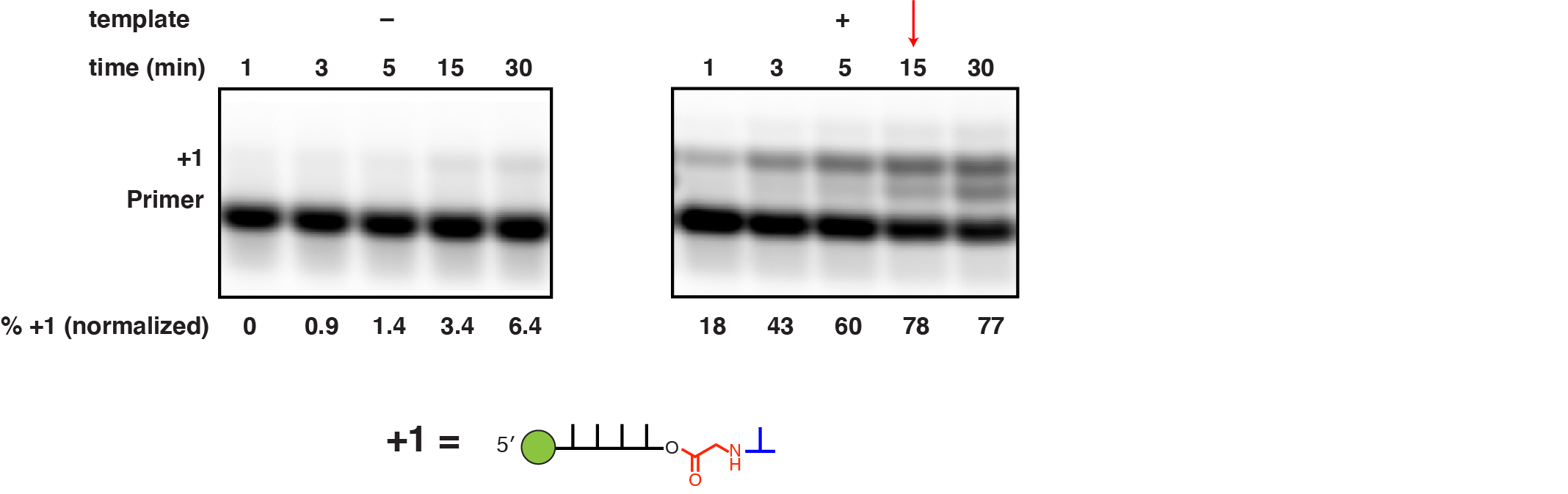

**Figure S6**

Off-template primer extension reaction of aminoacylated RNA 10mer primer.

A mixture of aminoacylated and non-aminoacylated fluorescently-labeled RNA 10mer primer was subjected to primer extension (200 mM HEPES pH 8.0, 20 mM C*C dimer, 2.5 mM MgCl_2_) with or without adding the RNA template. Primer extension products were monitored by denaturing 20% Urea-PAGE. The red arrow indicates the point where the templated reaction reached completion. The percentage of the +1 product was normalized to acylation efficiency that was separately measured to be 34% using acidic Urea-PAGE.

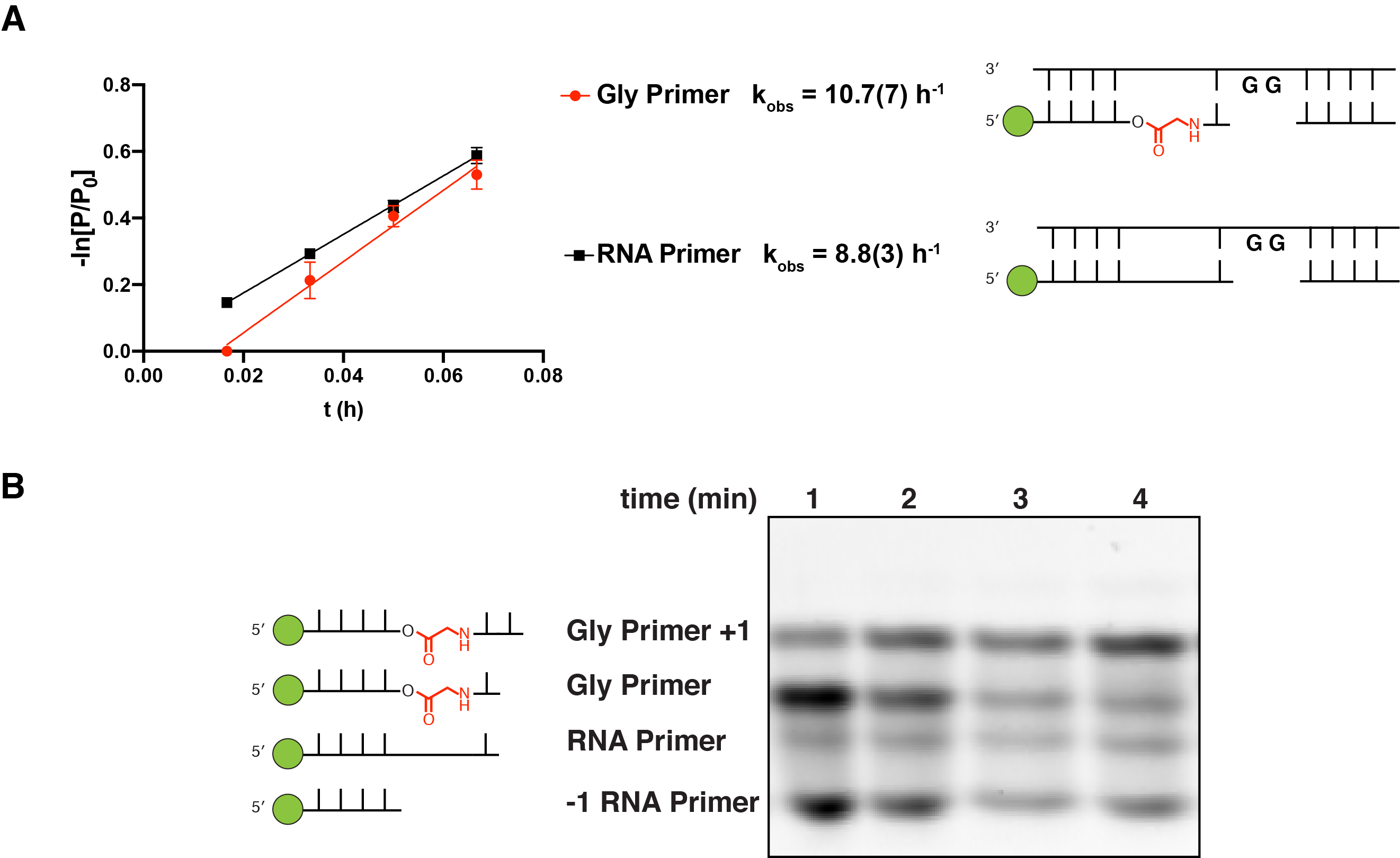

**Figure S7**

Kinetics of primer extension after the glycine linkage in the primer.

A. First-order kinetics of the RNA primer and glycine-linked primer extension reactions. Shown, on the left, is the plot of the negative natural logarithm of the ratio of primer (P) to initial primer (P_0_) versus time of reaction. Gly primer reaction data are colored red. Pure RNA primer reaction data are colored black. The slope of the linear fits corresponded to k_obs_ for the two reactions. Shown, on the right, is the schematic representation of the Gly primer and the pure RNA primer aligned to the template before reaction initiation. All reactions were performed at 200 mM HEPES pH 8.0, 50 mM MgCl_2_, with 20 mM C*C dimer. Values are reported as the mean +/− standard deviation from triplicate experiments.
B. Primer extension reactions were monitored by 20% denaturing Urea-PAGE. Shown gel is from the reaction of glycine-linked (Gly) primer (RNA reaction not shown). In addition to the Gly primer and its +1 extension product bands, the -1 RNA primer and its +1 extension product (RNA primer) bands were also observed. The -1 RNA primer is present due to the incomplete purification of the Gly primer (see Figures S2A and S2C for the synthesis and purification of Gly primer **1**). The data in A was obtained by monitoring the disappearance of the band labeled ‘Gly Primer’ (see **Kinetic analysis of primer extension reactions**).

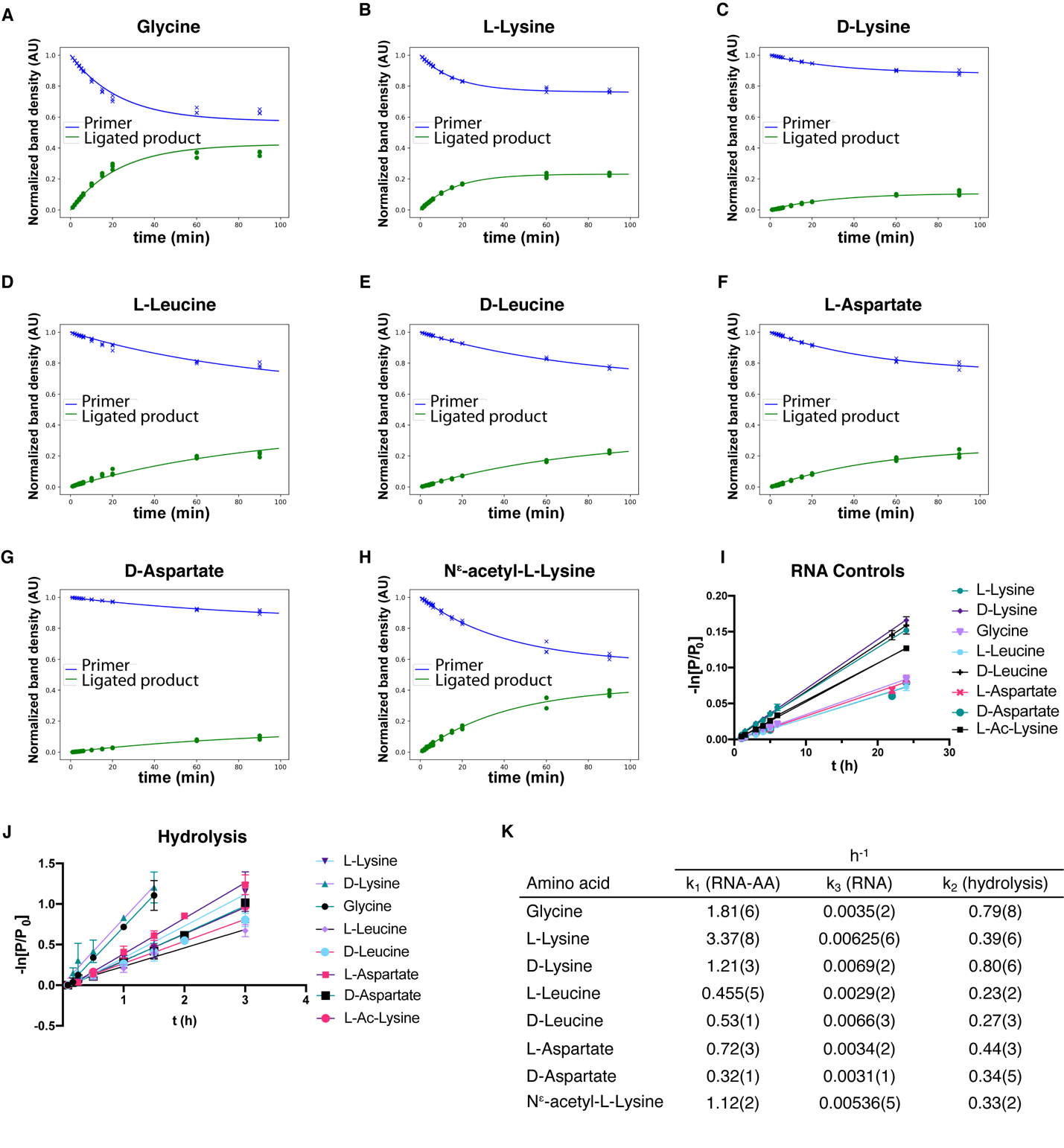

**Figure S8**

Kinetic analysis of ligation from 3′ hydroxyl and 2′(3′) aminoacyl terminated primers.

A.-H. Plots showing the normalized band intensities for the primer band, which includes both the primer-amino acid and primer species, (blue) and the ligated product (green) over the course of a ligation reaction with 2-methylimidazole activated 10mer. Non-linear fits were obtained using the kinetic model described in the Materials and Methods section.
I. RNA primer control ligation reactions in the presence of dinitrobenzyl esters of each of the tested amino acids. Shown is the plot of the negative natural logarithm of the ratio of primer (P) to initial primer (P_0_) versus time of ligation for a pure RNA primer. The slope of the linear fits corresponded to k_obs_ for RNA reactions that we defined as k_3_ in the kinetic model.
J. Primer-amino acid hydrolysis reactions. Shown is the plot of the negative natural logarithm of the ratio of primer-amino acid (P) to initial primer-amino acid (P_0_) versus time of reaction for the aminoacylated primer. The slope of the linear fits corresponded to k_obs_ for primer-amino acid hydrolysis that we defined as k_2_ in the kinetic model.
K. The tabulated summary of the obtained kinetic parameters from A-I plots. The values are reported as the mean +/− standard deviation from triplicate experiments.

Reaction conditions: 2.5 mM MgCl_2_, 200 mM HEPES pH 8.0, 2.5 µM primer, 3.75 µM template, 10 µM ligator. For the hydrolysis experiment 10 µM of unactivated ligator was used instead of the 2-metylimidazole activated ligator. Values are reported as the mean +/− standard deviation from triplicate experiments.

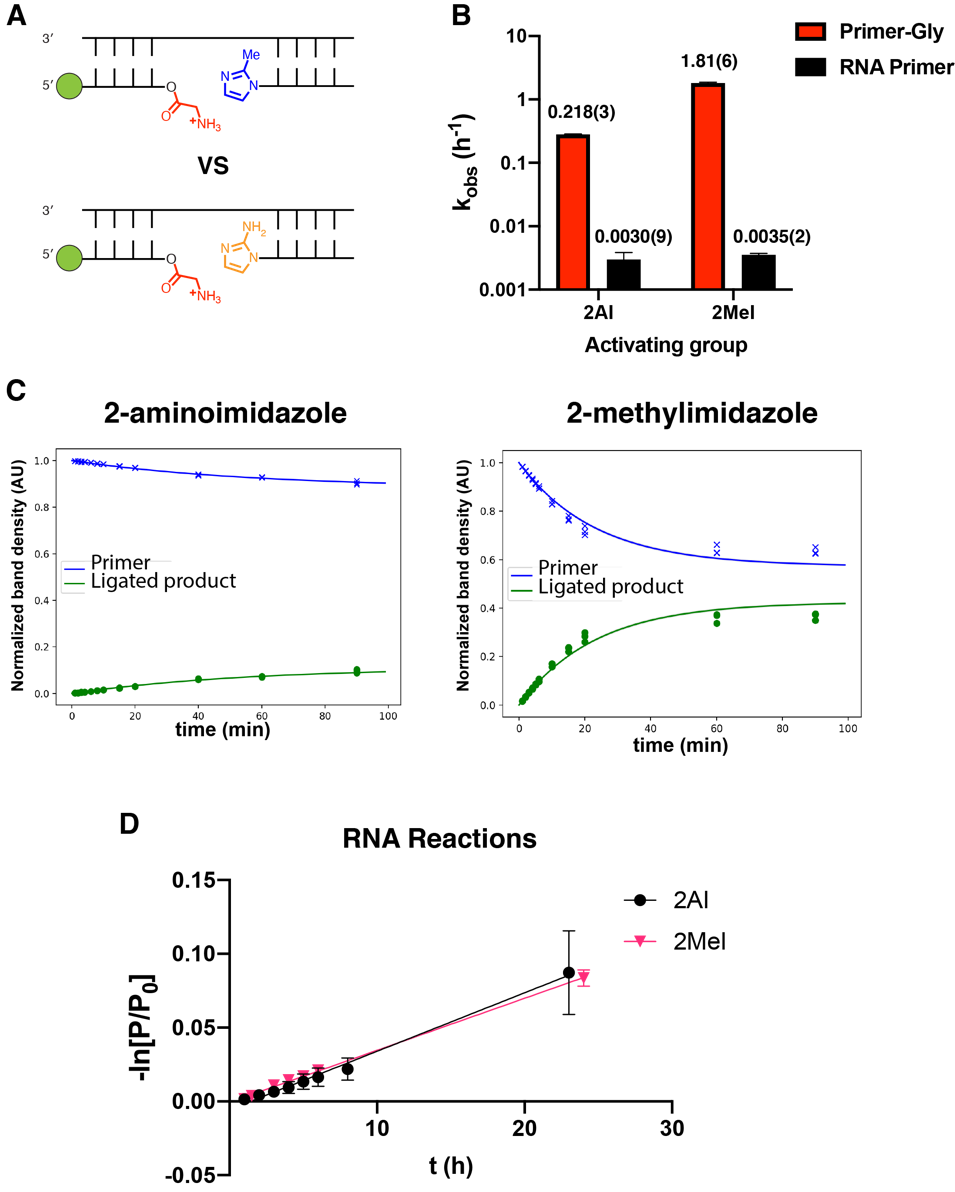

**Figure S9**

Kinetics of ligation reactions using 2-aminoimidazole or 2-methylimidazole activated 10mers.

A. Schematic of the reactions that were compared. Aminoacylated fluorescently-labeled 12mer RNA primers are shown. Pure non-aminoacylated 12mer RNA reactions were compared as well but are not shown in the diagram.
B. Observed rate constants for the ligation reactions of aminoacylated or non-aminoacylated 12mer RNA with either 2-aminoimidazole or 2-methylimidazole activated 10mers. Reaction conditions: 2.5 mM MgCl_2_, 200 mM HEPES pH 8.0, 2.5 µM primer, 3.75 µM template, 10 µM activated ligator. Values are reported as the mean +/− standard deviation from triplicate experiments.
C. Plots showing the normalized band intensities for the primer band, which includes both the primer-gly and primer species, (blue) and the ligated product (green) over the course of a ligation reaction with 2-aminoimidazole (left) and 2-methylimidazole (right) activated oligonucleotides. Fits are obtained using the kinetic model described in the Materials and Methods section.
D. Pure RNA ligation reaction. Plot of the negative natural logarithm of the ratio of primer (P) to initial primer (P_0_) versus time of ligation for a pure RNA primer with 2-aminoimidazole (black) and 2-methylimidazole (pink) activated oligonucleotides. The slope of the linear fits corresponds to k_obs_ for RNA.

Hydrolysis data for primer-gly can be found in Figure S8.

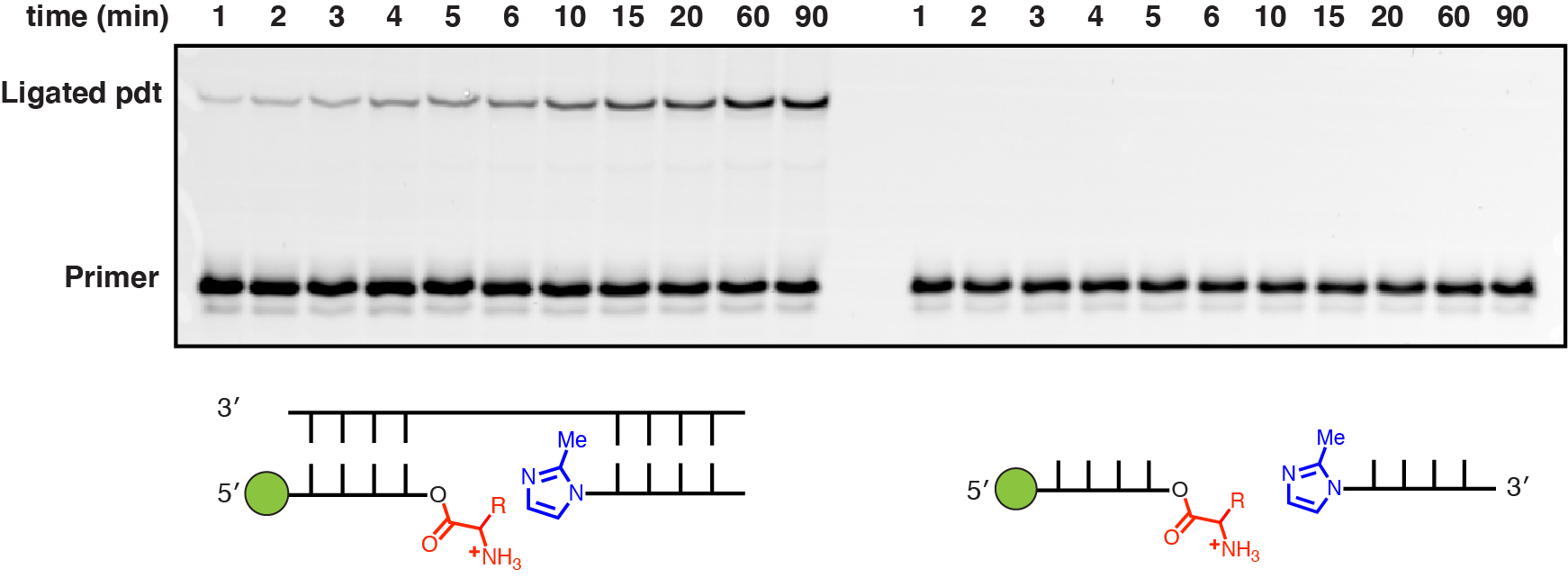

**Figure S10**

Off-template ligation reaction with aminoacylated fluorescently-labeled 12mer RNA primer and 2-methylimidazole activated 10mer. Reaction conditions were 2.5 mM MgCl_2_, 200 mM HEPES pH 8.0, 2.5 µM primer, 10 µM activated ligator with (left) or without (right) 3.75 µM RNA template.

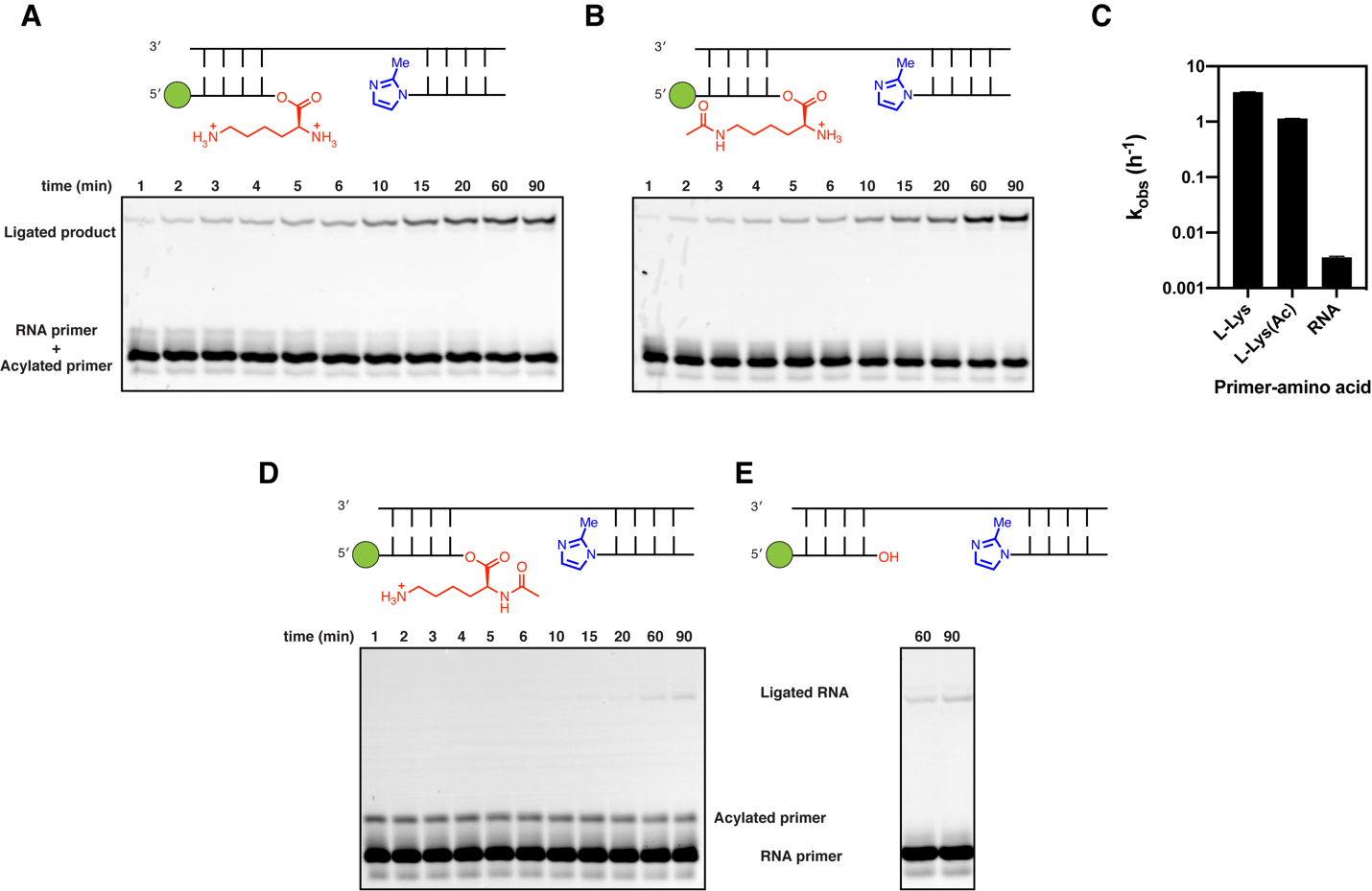

**Figure S11**

Ligation reactions with L-lysine and *N*^ε,α^-acetyl-L-lysine aminoacylated fluorescently-labeled 12mer RNA primer.

A.-B. A mixture of L-lysine or *N*^ε^*-*acetyl-L-lysine aminoacylated and non-aminoacylated fluorescently-labeled 12mer RNA primer was subjected to ligation conditions and reaction progress was followed by 20% denaturing Urea-PAGE. The starting mixture of primers is indistinguishable by gel. The ligated product appears after 1 minute, consistent with the phosphoramidate bond formation.
C. A bar plot summarizing the rates of reactions shown in panels A and B using the kinetic model described in **Materials and Methods**. Reaction rates for hydrolysis and RNA controls for each of the two reactions can be found in Figure S8.
D. A mixture of *N*^α^-acetyl-L-lysine aminoacylated and non-aminoacylated fluorescently-labeled 12mer RNA primer was subjected to ligation conditions and reaction progress was followed by 20% denaturing Urea-PAGE. The starting mixture of primers is distinguishable by gel, presumably due to the greater stability of *N*$\alpha$-acetyl-L-lysine compared to L-lysine aminoacyl ester. Ligated product only appears after 60 minutes and it likely corresponds to the background RNA ligation shown in C.
E. A pure RNA 12mer primer was subjected to typical ligation conditions and reaction progress was followed by 20% denaturing Urea-PAGE. Ligated product is only visible after 60 minutes.

Reaction conditions: 2.5 mM MgCl_2_, 200 mM HEPES pH 8.0, 2.5 µM primer, 3.75 µM template, 10 µM activated ligator.

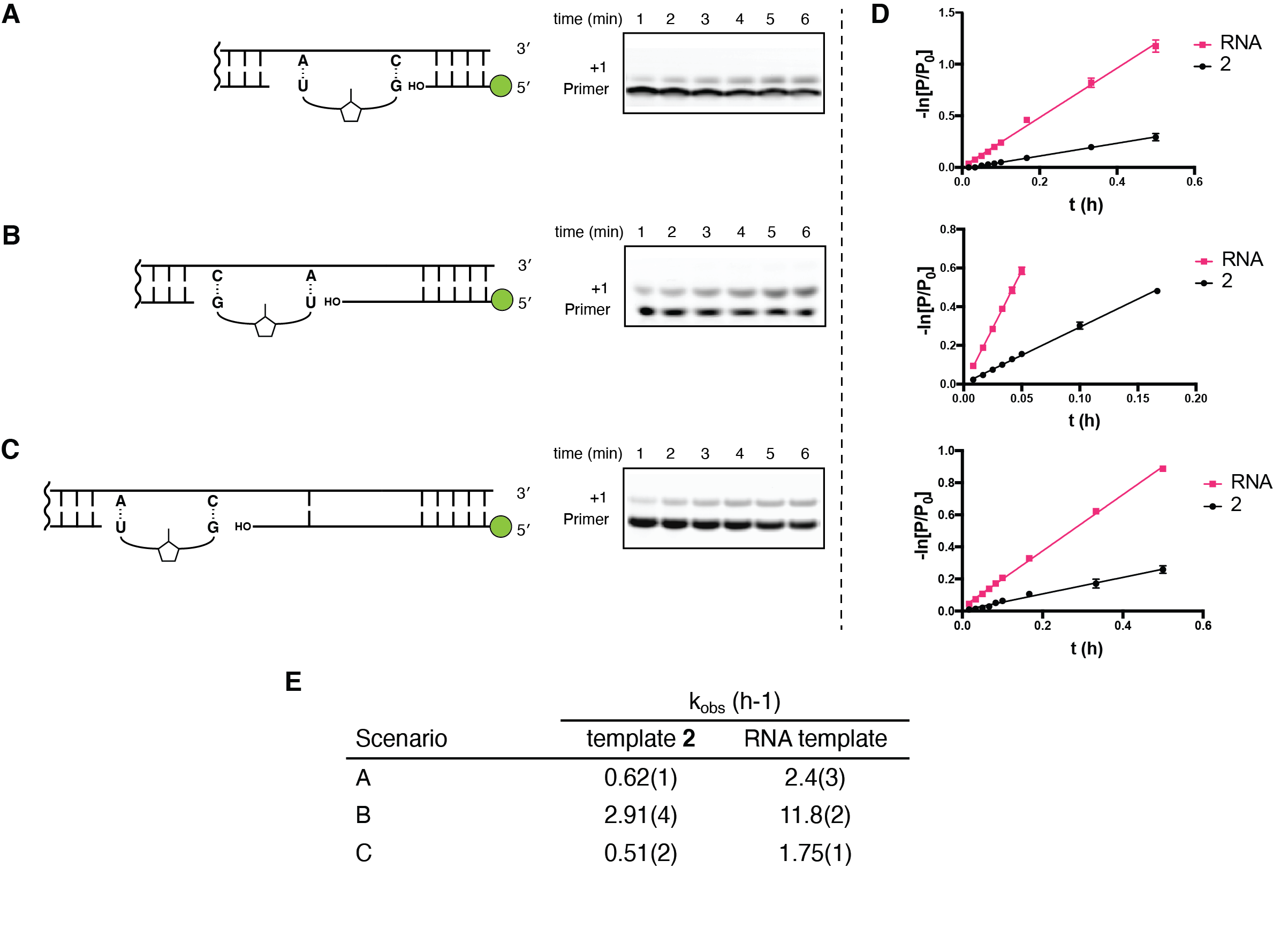

**Figure S12**

A.-C. Kinetic analysis of nonenzymatic primer extension across an RNA template without the amino acid bridge. Left: Schematic representation of the primer-template duplexes analyzed, showing the binding site for the G*U dimer in each case. Right: The time course of primer extension was monitored using polyacrylamide gel electrophoresis. All reactions were performed at 200 mM HEPES pH 8.0, 100 mM MgCl_2_, with 20 mM G*U dinucleotide.
D. First-order kinetic plots of primer extension reactions across a template containing a glycine bridge **2** (black) or RNA template without the bridge (pink). k_obs_ for each reaction was obtained from the slope of the linear fit of ln-transformed percentages of remaining primer over the course of reaction. Plots are aligned to correspond to the reactions in A-C.
E. Summary of obtained kinetic parameters from D. Values are reported as the mean +/− standard deviation from triplicate experiments.

**Supplementary Tables**

**Table S1:** Sequences used in this work. Nucleotides in magenta are deoxyribonucleotides, while all others are ribonucleotides.

| **Name** | **Sequence** | **Use** |
| --- | --- | --- |
| dFx Flexizyme | 5′-GGAUCGAAAGAUUUCCGCAUCCCC  GAAAGGGUACAUGGCGUUAGGU | Figure 2, 3, 4, S1, S2A |
| dFx Flexizyme M2 | 5′-GGAUCGAAAGAUUUCCGCAUCCCC  GAAAGGGUACAUGGCGUUAGUU | Figures 5, 6, S2B, S8, S9, S10, S11 |
| FX3 primer | 5′FAM-AGAGAAGCCA | Figures 2, 3, 4, S1, S2A, S5, S6 |
| FX_T2 template | 5′-UAAUCCAAGGUGGCUUCUCU | Figures 2, 3, 4 |
| FX_S2 sandwich | 5′-UUGGAUUA | Figures 2, 3, 4 |
| LXP1F primer | 5′FAM-AGAGAGCAGACA | Figures 5, S8, S9, S10, S11 |
| NP Lig T template | 5′-AGCUGCCGGGUGUCUGCUCUCU | Figures 5, S3, S8, S9, S10, S11; Table 1 (ds **2**) |
| Ligator1 | 5′-2MeImid (or 2AminoImid)CCCGGCAGCU | Figures 5, S2B, S8, S9, S10, S11 |
| LXP2F primer | 5′FAM-AGAGAAGAGAGCAGACA | Figures 6, S2B |
| NP DNA T template | 5′-AGCTGCCGGGTGTCTGCTCTCT | Figure S2B |
| FX3+1 primer | 5′FAM-AGAGAAGCCAC | Figure S7 |
| FX3+gly1 (**1**) primer | 5′FAM-AGAGAAGCCAglyC | Figures S2A, S3, S7; Table 1 |
| FX_T2 hybrid template | 5′-TAATCCAAGGUGGCTTCTCT | Figure S2A |
| FX_T5 template | 5′-UAAUCCAGGGUGGCUUCUCU | Figure S7; Table 1 (ds **1**) |
| FX_S5 sandwich | 5′-UGGAUUA | Figure S7 |
| NP FX1 primer | 5′FAM-AGCUGCCGG | Figure 6A, S12A |
| NP FX1 sandwich | 5′-GUCUGCUCUCU | Figure 6A, S12A |
| NP FX2 primer | 5′FAM-AGCUGCCGGG | Figure 6B, S12B |
| NP FX2 sandwich | 5′-UCUGCUCUCU | Figure 6B, S12B |
| NP FX3 primer | 5′FAM-AGCUGCCGGGU | Figure 6C, S12C |
| NP FX3 sandwich | 5′-CUGCUCUCU | Figure 6C, S12C |
| NP RNA T template | 5′-AGAGAGCAGACACCCGGCAGCU | RNA control template for Figure 6 |
| NP template (**2**) | 5′-AGAGAGCAGACAglyCCCGGCAGCU | Figure 6 template, S3, S4, S12; Table 1 |

**Characterization of G*U dinucleotide and 3,5-dinitrobenzyl esters of amino acids**

G*U 2-aminoimidazolium dinucleotide

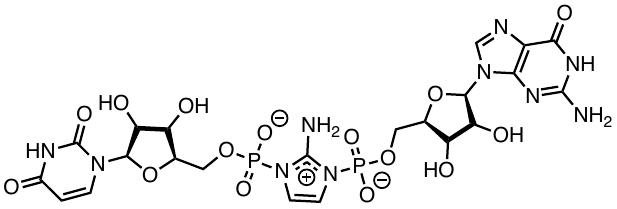

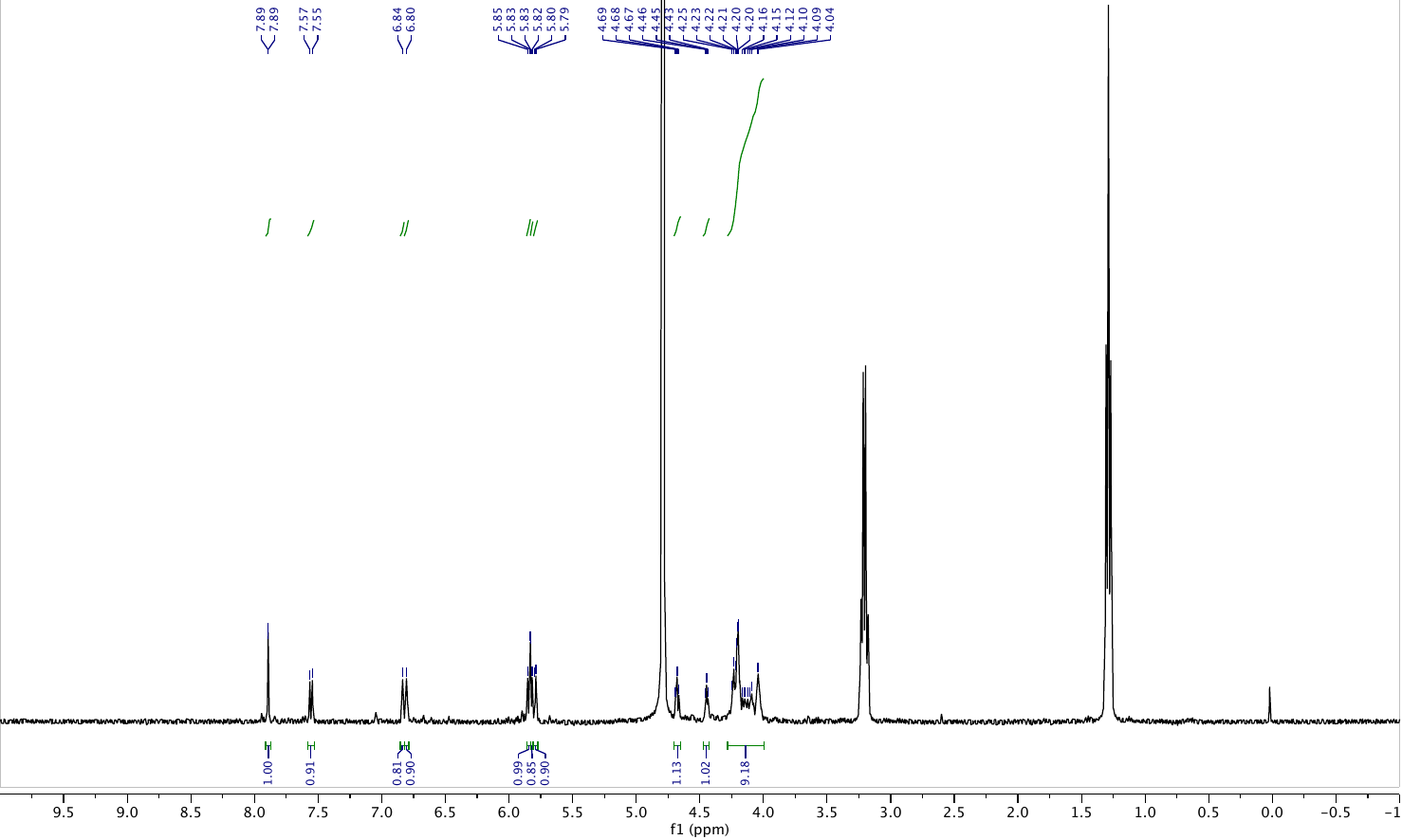

^1^H NMR (400 MHz, D_2_O) δ 7.89 (d, *J* = 1.3 Hz, 1H), 7.56 (d, *J* = 8.4 Hz, 1H), 6.84 (s, 1H), 6.80 (s, 1H), 5.84 (d, *J* = 8.3 Hz, 1H), 5.82 (d, *J* = 5.3 Hz, 1H), 5.79 (d, *J* = 4.3 Hz, 1H), 4.68 (t, *J* = 5.3 Hz, 1H), 4.45 (t, *J* = 4.7 Hz, 1H), 4.28 – 4.00 (m, 9H); peaks corresponding to residual TEAB were observed at 3.21 ppm and 1.29 ppm. LRMS (ESI- ion trap): calc for [C_22_H_27_N_10_O_15_P_2_]^–^ 733.11, found 733.2. HRMS (ESI-TOF): calc for [C_22_H_27_N_10_O_15_P_2_]^–^ 733.1138, found 733.1141.

G*U 2-aminoimidazolium dinucleotide

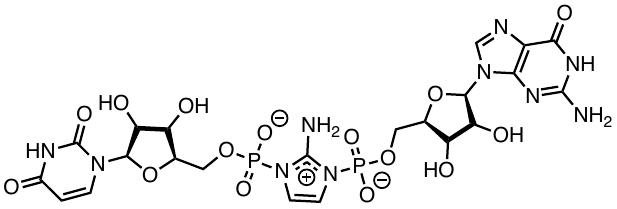

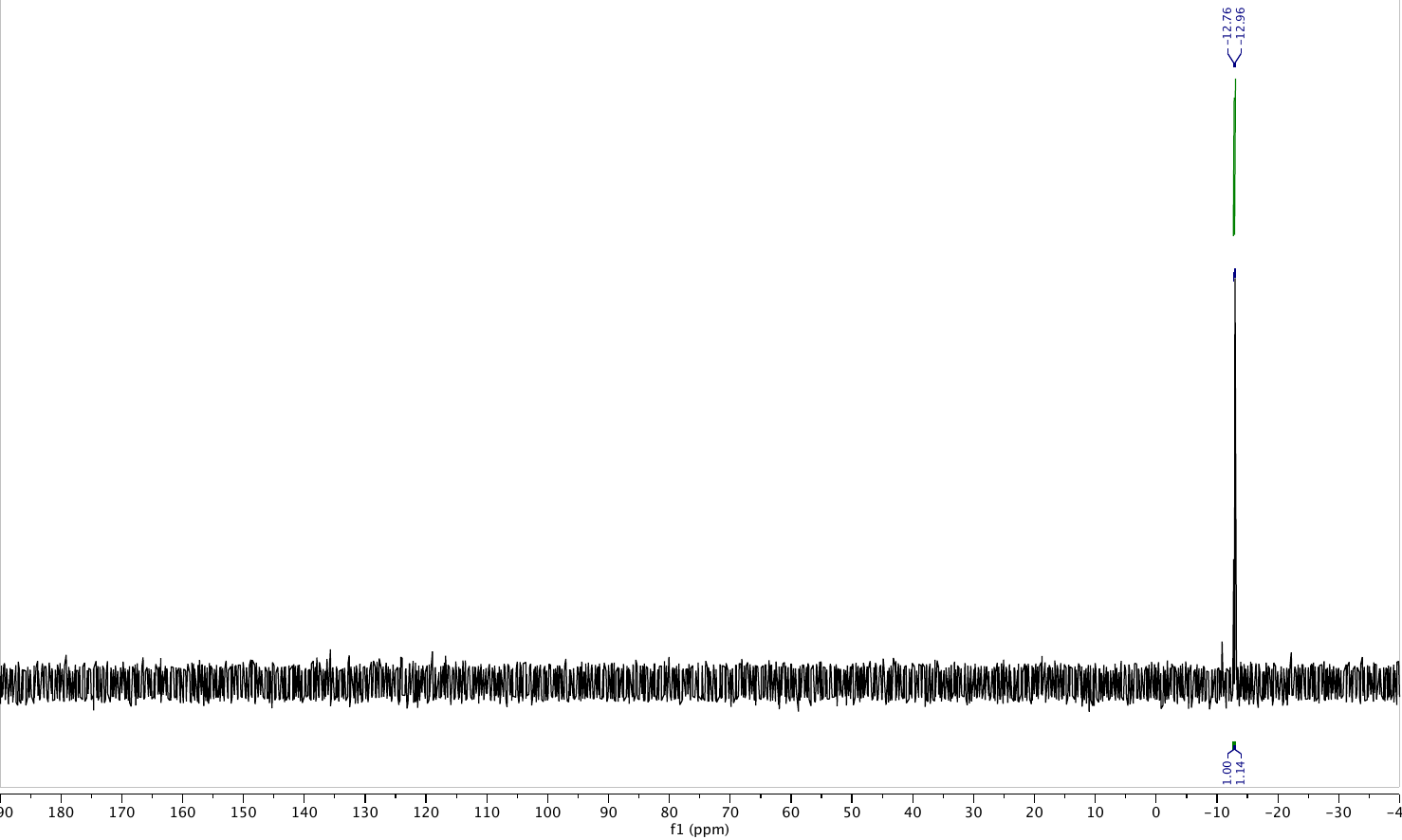

^31^P NMR (162 MHz, D_2_O) δ -12.76, -12.96.

(*S*)-3-amino-4-((3,5-dinitrobenzyl)oxy)-4-oxobutanoic acid or L-aspartate-DBE

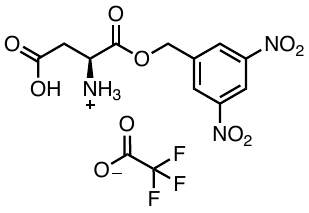

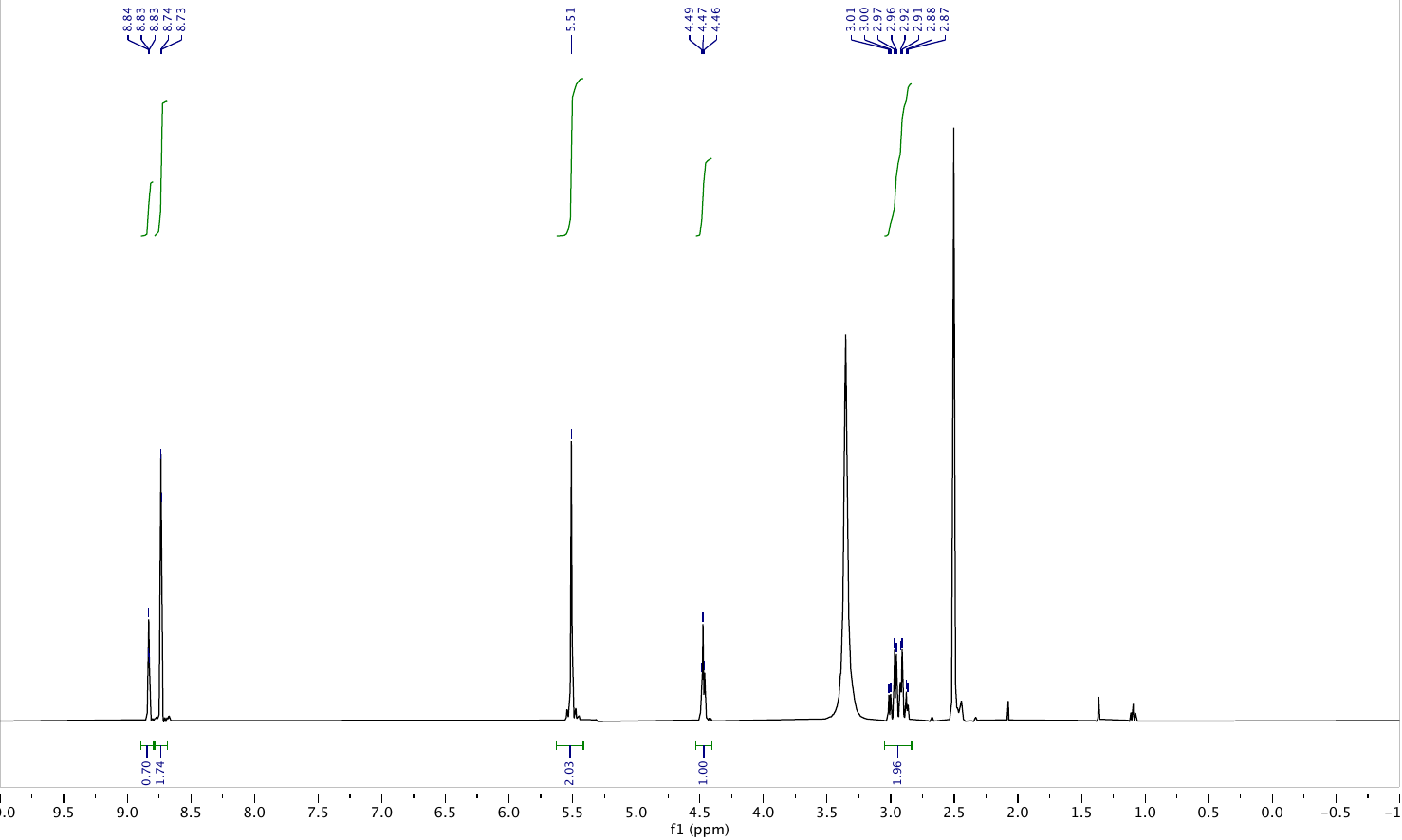

^1^H NMR (400 MHz, dmso) δ 8.83 (t, *J* = 2.1 Hz, 1H), 8.73 (d, *J* = 2.1 Hz, 2H), 5.51 (s, 2H), 4.47 (t, *J* = 5.1 Hz, 1H), 2.94 (qd, *J* = 18.0, 5.1 Hz, 2H). LRMS (ESI- ion trap): calc for [C_11_H_11_N_3_O_8_ + (H^+^)]^+^ 314.06, found 314.0. HRMS (ESI-TOF): calc for [C_11_H_11_N_3_O_8_ + (H^+^)]^+^ 314.06244, found 314.0642.

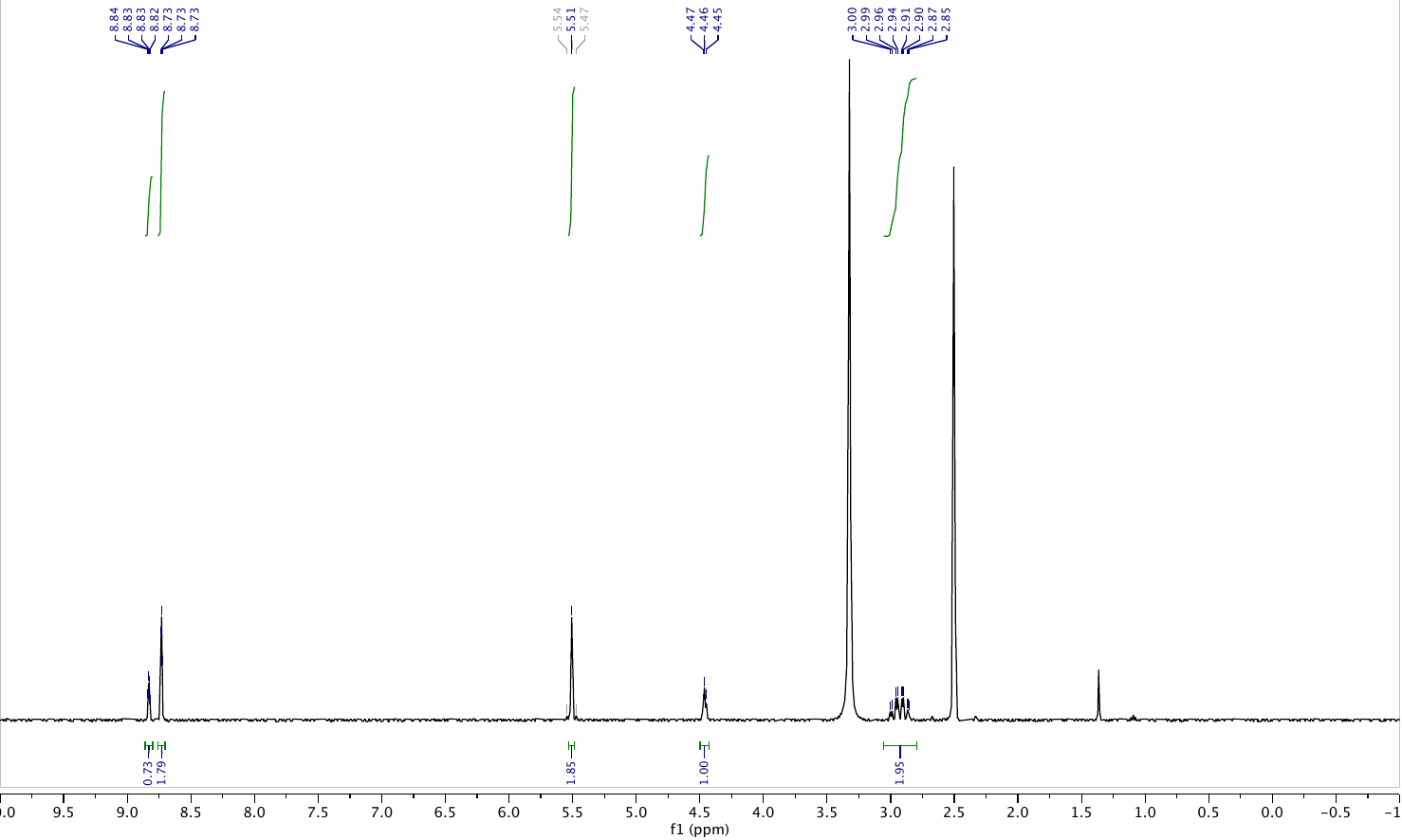
(*R*)-3-amino-4-((3,5-dinitrobenzyl)oxy)-4-oxobutanoic acid
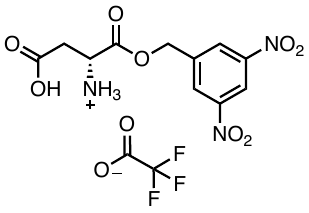
or D-aspartate-DBE

^1^H NMR (400 MHz, dmso) δ 8.83 (q, *J* = 2.0 Hz, 1H), 8.73 (d, *J* = 1.9 Hz, 2H), 5.51 (s, 2H), 4.46 (t, *J* = 5.1 Hz, 1H), 2.93 (qd, *J* = 17.8, 4.9 Hz, 2H). LRMS (ESI- ion trap): calc for [C_11_H_11_N_3_O_8_ + (H^+^)]^+^ 314.06, found 314.0. HRMS (ESI-TOF): calc for [C_11_H_11_N_3_O_8_ + (H^+^)]^+^ 314.06244, found 314.0639.

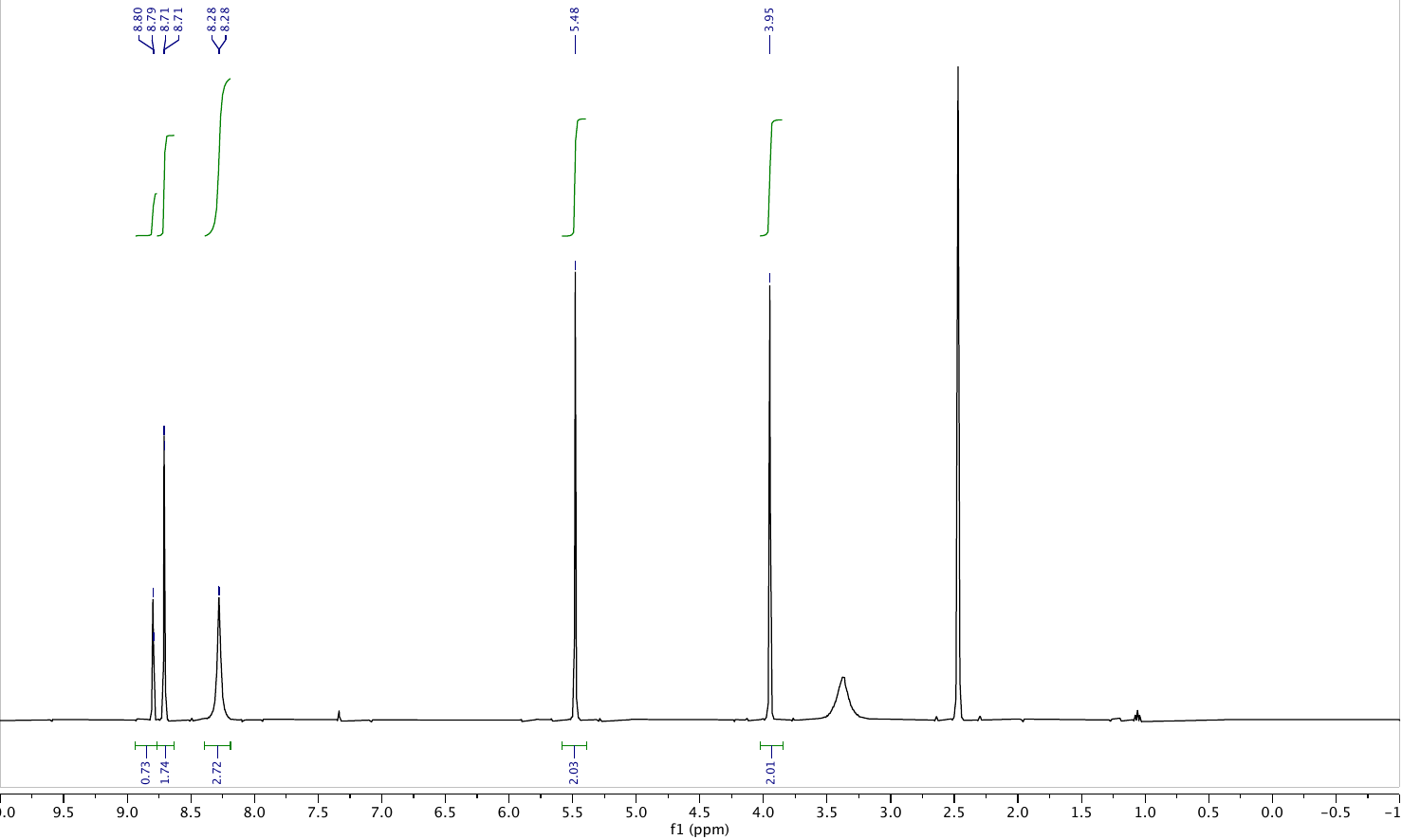
3,5-dinitrobenzyl glycinate
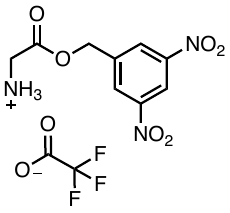
or glycine-DBE

^1^H NMR (400 MHz, dmso) δ 8.80 (d, *J* = 2.2 Hz, 1H), 8.71 (d, *J* = 2.1 Hz, 2H), 8.39 – 8.19 (m, 3H), 5.48 (s, 2H), 3.95 (s, 2H). LRMS (ESI- ion trap): calc for [C_9_H_9_N_3_O_6_ + (H^+^)]^+^ 256.06, found 255.9. HRMS (ESI-TOF): calc for [C_9_H_9_N_3_O_6_ + (H^+^)]^+^ 256.05696, found 256.0582.

3,5-dinitrobenzyl *L*-leucinate
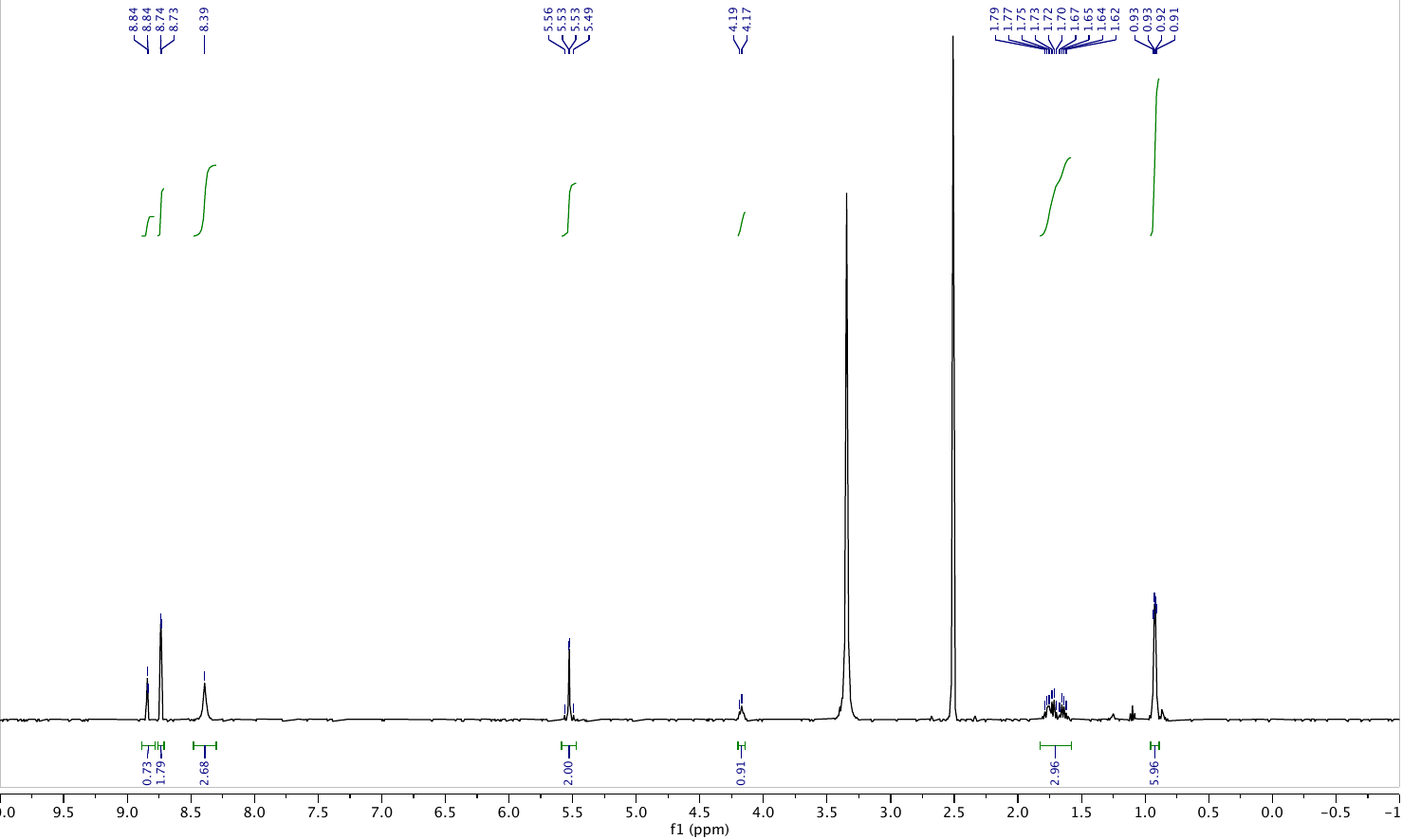

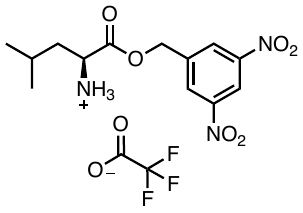
or L-leucine-DBE

^1^H NMR (400 MHz, dmso) δ 8.84 (d, *J* = 2.1 Hz, 1H), 8.74 (d, *J* = 2.1 Hz, 2H), 8.39 (s, 3H), 5.53 (d, *J* = 1.9 Hz, 2H), 4.18 (d, *J* = 7.2 Hz, 1H), 1.83 – 1.58 (m, 3H), 0.92 (dd, *J* = 6.3, 3.0 Hz, 6H). LRMS (ESI- ion trap): calc for [C_13_H_17_N_3_O_6_ + (H^+^)]^+^ 312.12, found 312.1. HRMS (ESI-TOF): calc for [C_13_H_17_N_3_O_6_ + (H^+^)]^+^ 312.11956, found 312.1218.

3,5-dinitrobenzyl *D*-leucinate
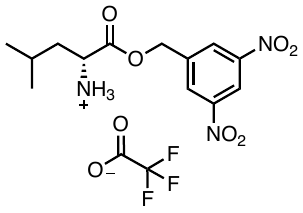

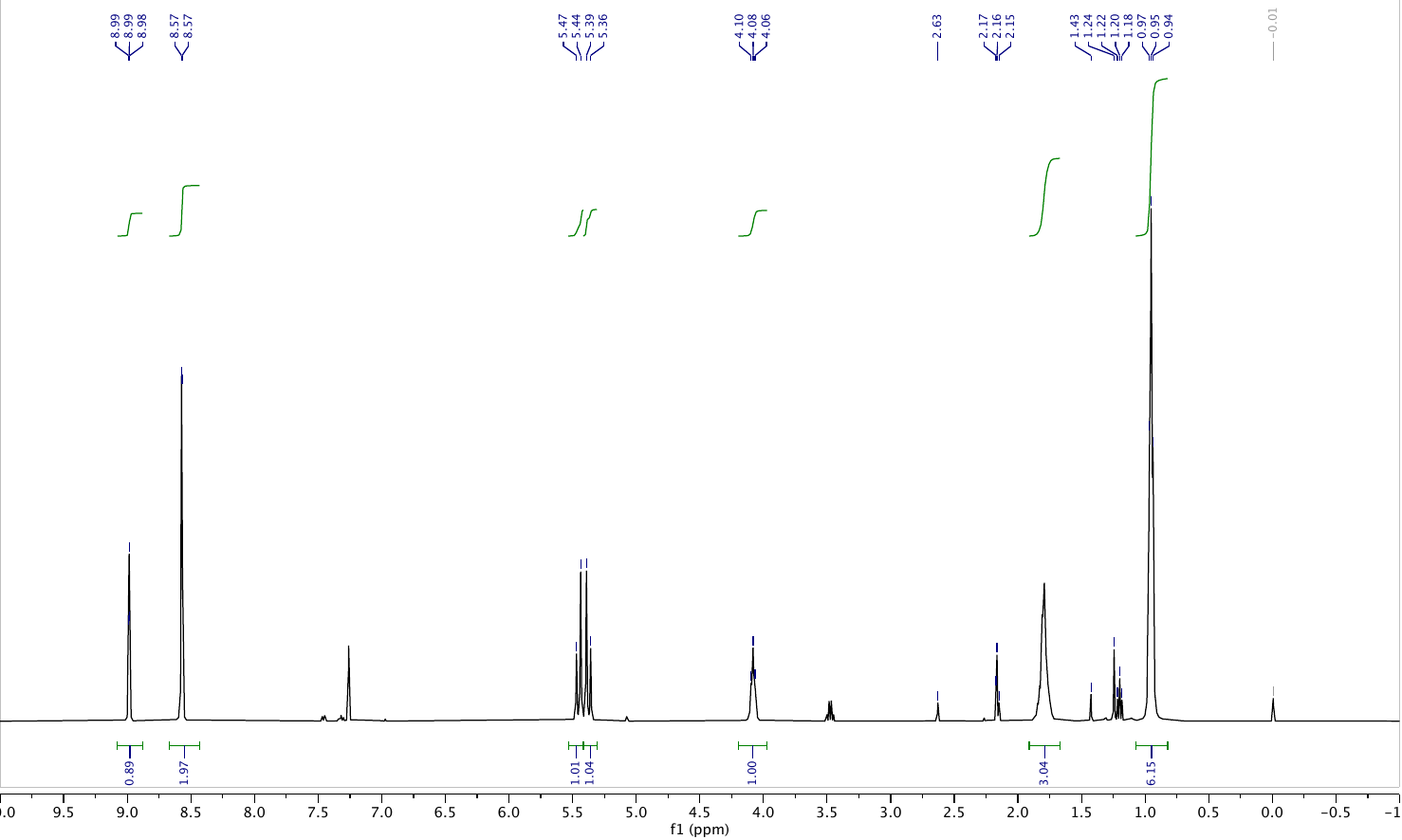
or D-leucine-DBE

^1^H NMR (400 MHz, cdcl_3_) δ 8.99 (t, *J* = 2.1 Hz, 1H), 8.57 (d, *J* = 2.1 Hz, 2H), 5.45 (d, *J* = 13.2 Hz, 1H), 5.37 (d, *J* = 13.2 Hz, 1H), 4.20 – 3.97 (m, 1H), 1.79 (p, *J* = 7.7, 6.7 Hz, 3H), 0.95 (t, *J* = 6.0 Hz, 6H). LRMS (ESI- ion trap): calc for [C_13_H_17_N_3_O_6_ + (H^+^)]^+^ 312.12, found 312.1. HRMS (ESI-TOF): calc for [C_13_H_17_N_3_O_6_ + (H^+^)]^+^ 312.11956, found 312.1210.

­­

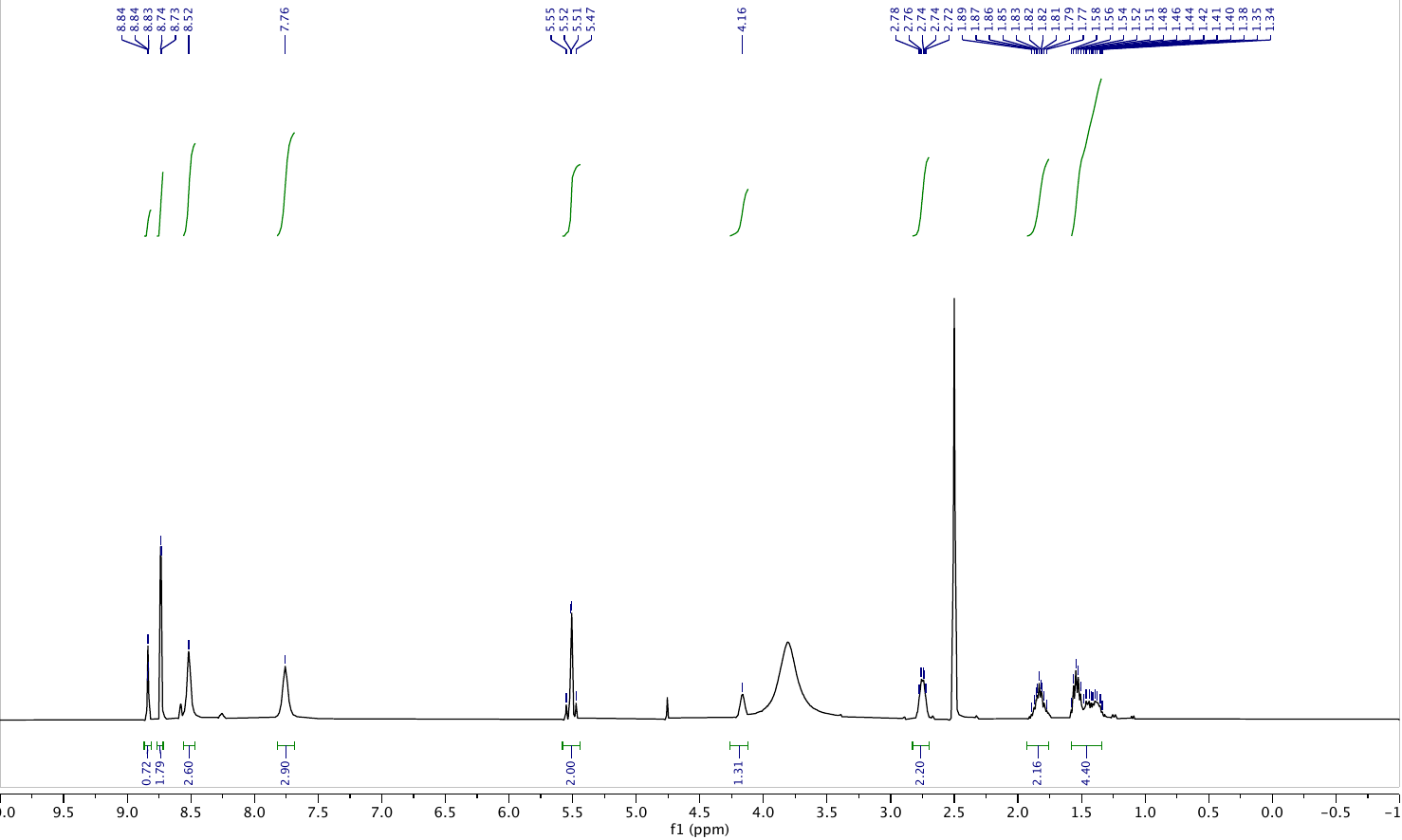

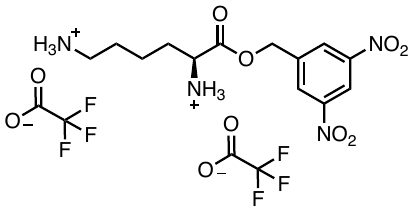
 3,5-dinitrobenzyl *L*-lysinate or L-lysine-DBE

^1^H NMR (400 MHz, dmso) δ 8.84 (t, *J* = 2.2 Hz, 1H), 8.74 (d, *J* = 2.1 Hz, 2H), 8.52 (s, 3H), 7.76 (s, 3H), 5.58 – 5.44 (m, 2H), 4.16 (s, 1H), 2.75 (q, *J* = 6.8 Hz, 2H), 1.83 (qd, *J* = 9.5, 6.1 Hz, 2H), 1.58 – 1.34 (m, 4H). LRMS (ESI- ion trap): calc for [C_13_H_18_N_4_O_6_ + (H^+^)]^+^ 327.13, found 327.1. HRMS (ESI-TOF): calc for [C_13_H_18_N_4_O_6_ + (H^+^)]^+^ 327.13046, found 327.1312.

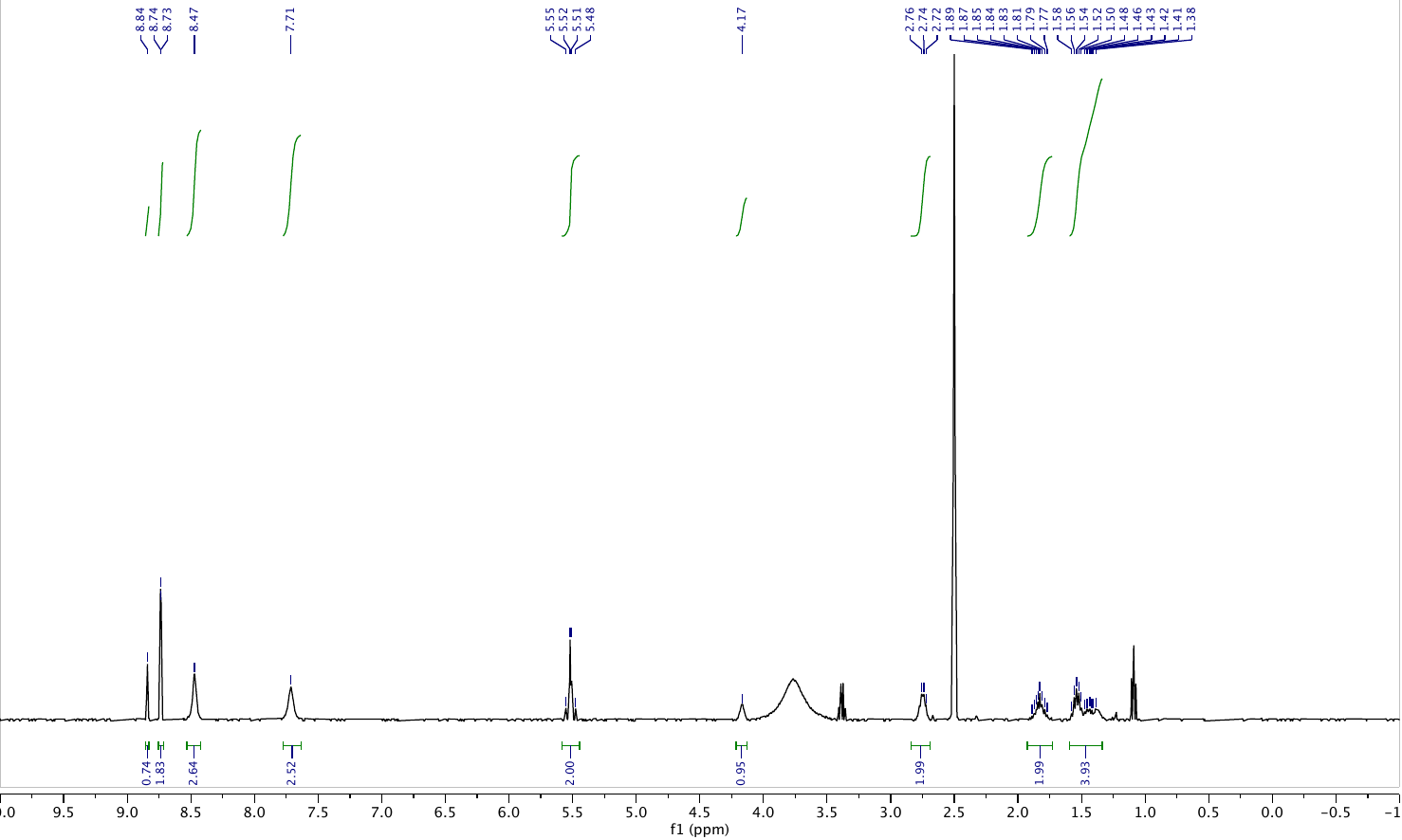
3,5-dinitrobenzyl *D*-lysinate
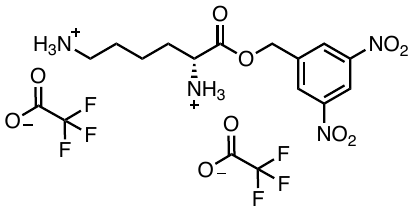
or D-lysine-DBE

^1^H NMR (400 MHz, dmso) δ 8.84 (s, 1H), 8.74 (d, *J* = 2.1 Hz, 2H), 8.47 (s, 3H), 7.71 (s, 3H), 5.59 – 5.44 (m, 2H), 4.17 (s, 1H), 2.84 – 2.69 (m, 2H), 1.83 (dh, *J* = 16.9, 7.8 Hz, 2H), 1.49 (ddt, *J* = 36.9, 20.7, 10.9 Hz, 4H). LRMS (ESI- ion trap): calc for [C_13_H_18_N_4_O_6_ + (H^+^)]^+^ 327.13, found 327.1. HRMS (ESI-TOF): calc for [C_13_H_18_N_4_O_6_ + (H^+^)]^+^ 327.13046, found 327.1329.

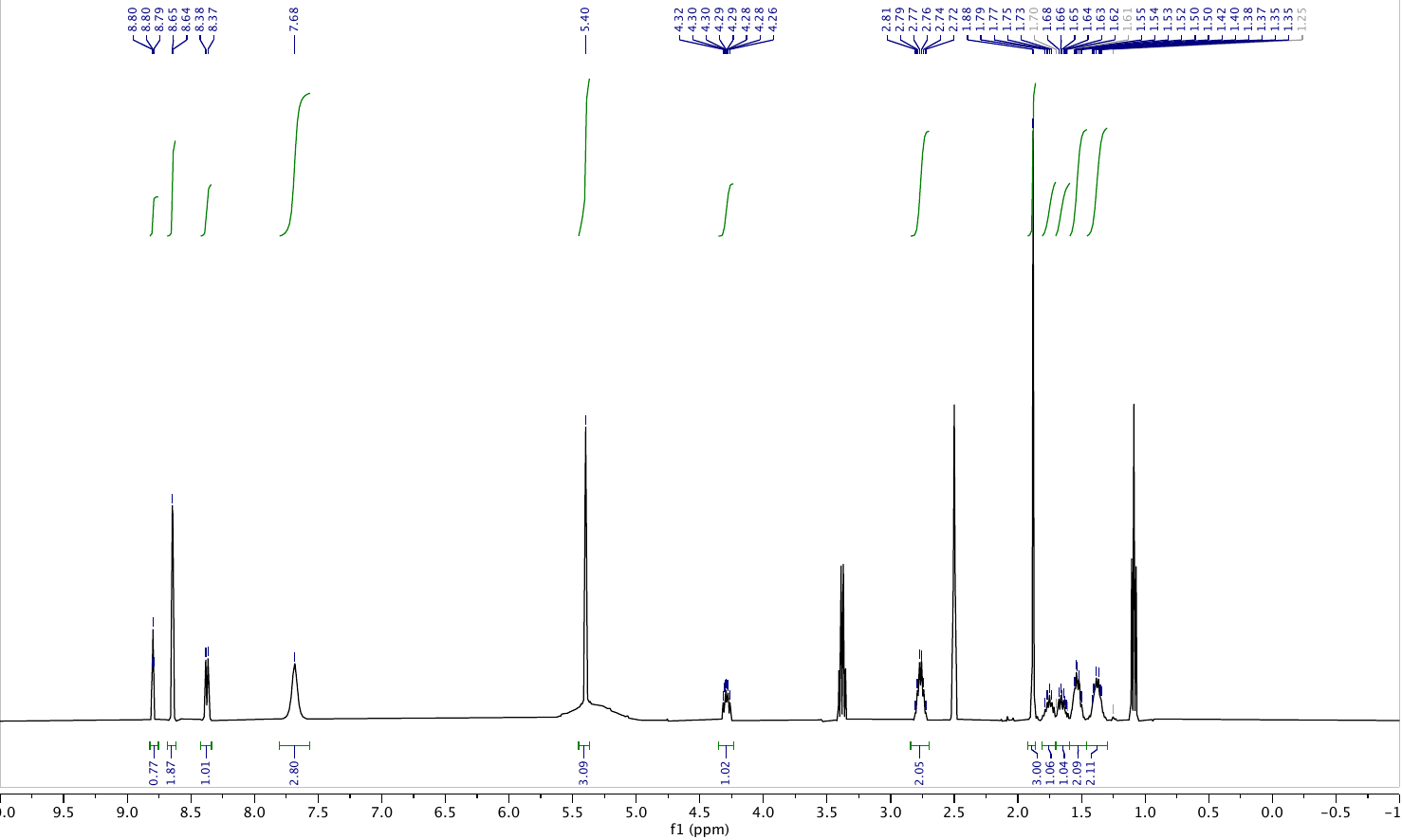
3,5-dinitrobenzyl acetyl-*L*-lysinate
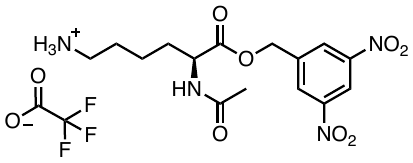
or L-acetyllysine-DBE

^1^H NMR (400 MHz, dmso) δ 8.80 (t, *J* = 2.1 Hz, 1H), 8.64 (d, *J* = 2.1 Hz, 2H), 8.37 (d, *J* = 7.2 Hz, 1H), 7.68 (s, 3H), 5.40 (s, 3H), 4.29 (ddd, *J* = 9.0, 7.1, 5.2 Hz, 1H), 2.76 (h, *J* = 6.1 Hz, 2H), 1.88 (s, 3H), 1.76 (dq, *J* = 13.7, 6.7 Hz, 1H), 1.70 – 1.59 (m, 1H), 1.52 (td, *J* = 8.6, 3.7 Hz, 2H), 1.38 (q, *J* = 6.7, 5.4 Hz, 2H). LRMS (ESI- ion trap): calc for [C_15_H_20_N_4_O_7_ + (H^+^)]^+^ 369.14, found 369.1. HRMS (ESI-TOF): calc for [C_15_H_20_N_4_O_7_ + (H^+^)]^+^ 369.14102, found 369.1430.
